# Turnip mosaic virus co-opts host RNA methylation machinery to orchestrate plant infection

**DOI:** 10.64898/2026.05.08.723783

**Authors:** Nicola Secco, Kashif Nawaz, Marilia Almeida-Trapp, Arsheed H. Sheikh, Heribert Hirt

**Author notes:** King Abdullah University of Science and Technology, Jeddah, Thuwal, Saudi Arabia (Nicola Secco).

## Abstract

N6-methyladenosine (m^6^A) is a key RNA modification that regulates transcript stability and translation. However, its function in plant viruses remains largely unclear. Here, we show that the positive-sense single-stranded +ssRNA Turnip mosaic virus (TuMV) relies on the host m^6^A machinery to support efficient infection. Our findings uncover a previously unrecognized nuclear phase in the TuMV life cycle, during which viral RNA undergoes extensive methylation by host enzymes. We identify a complex and non-canonical methylation landscape on the TuMV genome, where canonical DRACH motifs are embedded within clusters of additional virus-specific non-canonical m^6^A sites. Notably, we also detect the presence of another RNA modification, m^5^C (5-methylcytosine), in close proximity to m^6^A-marked regions. This coordinated methylation landscape appears to be critical for efficient viral polyprotein synthesis. In its absence, the virus displays aberrant methylation and reduced infectivity as observed in m^6^A writer and reader mutants. Based on this, we propose a “seeding” model in which initial m^6^A deposition at canonical sites nucleates the formation of a broader network of m^6^A and m^5^C marks, likely guided by RNA structure rather than sequence motifs alone.

## Introduction

Among all RNA modifications, N^6^-methyladenosine (m^6^A) is the most abundant and well-studied modification in eukaryotic cells. m^6^A plays a significant role in regulating various aspects of RNA metabolism, including splicing, stability, translation, and degradation of mRNAs. This modification is dynamically regulated by methyltransferases known as writers and demethylases known as erasers, while m^6^A sites are recognized by m^6^A - binding proteins known as readers. These methyltransferases can be grouped into distinct categories based on their targets, ranging from mRNA to rRNA and tRNA^1–4^. A multienzymatic complex with a mass of 1 MDa is the most prominent adenosine methyltransferase for mRNA and lncRNA in metazoan organisms. The methyltransferase METTL3 and its cofactor METTL14 comprise the complex catalytic core^5–7^. The orthologues of METTL14 and METTL3 in Arabidopsis are MTB and MTA, respectively^8,9^. MTA functions as an active methyltransferase enzyme and exhibits characteristics similar to its metazoan counterpart^9^. MTB, despite lacking methyltransferase activity in Arabidopsis, affects the localization and catalytic activity of MTA^8,10^. However, MTB may demonstrate catalytic activity in other plant species^11^. The MTA and MTB heterodimer stability is enhanced by FKBP12 Interacting Protein 37 (FIP37)^8,10^. Another protein in the complex is VIR1, a homolog of the sex-determining Drosophila VIRILIZER^12,13^, which serves as a scaffolding protein for the methyltransferase complex^8,10,14,15^. HAKAI, an E3-ubiquitin ligase, is also considered to be part of the methyltransferase complex. While the precise function of HAKAI in the methyltransferase complex remains uncertain, *hakai* mutants show significant reduction in m^6^A deposition^8,10,16,17^. Recent proteomic studies have expanded the known architecture of the Arabidopsis m^6^A writer complex by identifying HIZ1 and HIZ2 as HAKAI-interacting zinc finger proteins^18^. Despite significant structural divergence, HIZ2 appears to be a functional equivalent of the Drosophila Flacc and human ZC3H13. *hiz2* mutants exhibit a substantial reduction in global m^6^A levels, underscoring its essential role in the methylation process^18^. Conversely, HIZ1 has been proposed as a functional analogue to mammalian ZFP217, acting as a negative regulator or “fine-tuner” of methylation^18^. While m^6^A levels remain stable in *hiz1* knockout mutants, its overexpression leads to a significant decrease in m^6^A abundance and developmental phenotypes reminiscent of core writer complex mutants^18^.

Reader proteins contain YTH domains, which are characterized by three tryptophan residues that form a hydrophobic pocket^19^. 13 genes in Arabidopsis were identified and designated as EVOLUTIONARY CONSERVED C-TERMINAL REGION (ECTs) based on the presence of a YTH domain^20^. The majority of the ECT1-11 proteins are homologous to human YTHDFs and are likely localized in the cytoplasm. Conversely, ECT12/13 demonstrate a high degree of similarity to YTHDCs, suggesting that they are likely nuclear proteins^20^.

The ALKBH family were identified as primary erasers in animals, due to their nonheme Fe(II) ⍺ −ketoglutarate-dependent-dioxygenase-like (ALKB-like) domain^21^. In plants, 13 homologs of the ALKBH family have been identified^22,23^.

Another prominent methylation present in mRNA is m^5^C. Similar to m^6^A m, m^5^C modification is catalyzed by writer methyltransferases^24,25^. These act primarily as single enzymes rather than complexes. In planta, the most prominent m^5^C writers have been identified as homologs ofanimal NSUN2 (TRM4A and TRM4B in *Arabidopsis*)^26–28^. ^5^C methylations are recognized by a set of reader proteins, primarily identified as homologs of the animal *ALY* genes, which have been shown to influence RNA localization^29,30^. Other m^5^C readers have also been identified in animals, such as SRSF2 which is involved in pre-mRNA splicing, and YBX1 which acts as RNA stabilizer^24,31,32^. It remains unclear which enzymes act as m^5^C demethylases in plants; however, potential candidates can be found within the *ALKBH* gene family, specifically the *ALKBH1* homologs, given that they have demonstrated m^5^C demethylase activity in animals^33^. Another possibility is that m^5^C modifications are inactivated by conversion in to 5-hydroxymethylcytidine (hm^5^C) by the activity of TET2^34–36^.

Viruses are minimal biological entities that often lack the necessary components for their own replication and spread, making them heavily reliant on the host’s metabolism for successful replication. Host RNA methylation plays a crucial role in the metabolism of animal viruses^37–42^, and recent studies have also highlighted its importance for plant viruses^11,23,43,43–50^. For example, Alfalfa Mosaic Virus (AMV), a positive single-stranded RNA virus (+ssRNA), is targeted by thost methyltransferases. AMV methylations are detrimental for the virus, leading AMV to rely on the host demethylase ALKBH9B to remove the m^6^A marks^23,48,50–52^. Other +ssRNA viruses, such as Potato Virus Y (PVY), Plum Pox Virus (PPV), and Tobacco Mosaic Virus (TMV), cause a general reduction in host m^6^A levels after infection. Although the precise mechanism remains unclear, this m^6^A reduction is likely a viral strategy to prevent the methylation of viral RNAs (vRNAs)^44,45^. Another +ssRNA virus, Pepino Mosaic Virus (PepMV), has developed a unique mechanism to counteract the host cell’s m^6^A machinery by directing HAKAI to autophagic degradation^46,47^. Additionally, some +ssRNA viruses, predominantly flexiviruses and potyviruses, have ALKB-like domains in their viral proteins, which were likely acquired through horizontal gene transfer^44,53,54^. This emphasizes the importance of demethylases in these viral life cycles. In most reported cases of plant-virus interactions, m^6^A has a negative effect on viral replication and tropism. However, there are instances where viruses benefit from m^6^A. For example, Wheat Yellow Mosaic Virus (WYMV) interacts with the host protein MTB, which gets relocated to the cytosol and is likely incorporated into Viral Replication Centers (VRCs)^11,49^. Additionally, m^5^C methylation has been shown to be largely involved in the virus-host interactions of both plants and animals. Similar to m^6^A, m^5^C hyper- or hypomethylation can lead to different outcomes depending on the specific virus-host system^24,29,55,56^.

Among plant viruses, potyviruses represent the largest family, including some of the most threatening pathogens for crop production^57^. These viruses have a +ssRNA genome consisting of a single open reading frame (ORF), flanked by 5’ and 3’ Untranslated Regions (UTRs)^58–61^. The vRNAs are polyadenylated and have a 5’ cap in the form of a viral genome-linked protein (VPg). The ORF codes for a polyprotein that is subsequently proteolytically cleaved into functional viral proteins by the viral proteases P1, HC-Pro, and NIa. Additionally, a secondary protein, PN3-PIPO, is formed through transcriptional slippage^62^.

Among potyviruses, Turnip Mosaic Virus (TuMV) is capable of infecting a wide range of wild and domesticated plants, causing yield losses of up to 70% in infected crops^63,64^. Potyviruses have been found to both influence and be influenced by host methylations^65–68^. However, it remains unclear if these interactions are conserved across this viral family and across different host-virus interactions. In this study, we investigated the impact of methylation on TuMV in *Arabidopsis thaliana* and *Nicotiana benthamiana*, and how these modifications affect viral titer and protein synthesis.

## Materials and methods

### Media and bacterial growth conditions

Bacteria were grown in LB BROTH BASE (12780029, Invitrogen) or in solid LB AGAR (22700041, Invitrogen) with the appropriate selection marker. *E. coli*-related strains were incubated for 24 hours at 37 °C, while *A. tumefaciens* (GV3101) was incubated at 28 °C for 48 hours.

### Vectors and expression systems

Agrobacterium with TuMV plant expression vector was obtained from Prof. Magdy M.Mahfouz lab^129^. Briefly, Turnip Mosaic Virus (TuMV) (NCBI accession number NC_002509.2) cDNA was cloned into a plant expression vector (Gene Bank accession number KX665539.1). The TuMV cDNA was further modified by inserting a Soluble-modified Red-shifted GFP (GFP) between the *P1* and *HC-Pro* cistrons, resulting in a recombinant virus expression vector referred to as *p35S::TuMV-GFP*. To ensure proper viral polyprotein processing, an NIa cleavage site was added at the C-terminus of the GFP. Potyviral viral RNAs (vRNAs) are translated as a single polyprotein, which is subsequently processed into functional viral proteins by the cis and trans activity of the viral-encoded proteases^61,130,131^. As a result, GFP is cleaved from the polyprotein and primarily found as an independent soluble protein. Due to its integration into the viral cistron structure, GFP is translated along with the rest of the polyprotein, providing an effective tool to identify the tissues in which TuMV is metabolically active.

The plasmids *pGdL5::GW* and *pGdL3::GW* were synthesized by GenScript using the *pGreen dual Luc ORF sensor* (Addgene #55207) as a template. The original vector was modified by inverting the promoters and appending C-terminal purification tags to each reporter (Rluc-HA and Fluc-FLAG). Finally, a Gateway (GW) cloning cassette was inserted at either the 5’ end (*pGdL5::GW*) or the 3’ end (*pGdL3::GW*) of the Fluc reporter gene.

PCR cloning of gene loci, cistrons, and coding sequences (CDS) was performed using Phusion™ High-Fidelity DNA Polymerase (F530L, Thermo Fisher) following the manufacturer’s specifications for primer design and reaction conditions. PCR products were screened through gel electrophoresis for verification.

*MTA* CDS (*MTA*) was isolated from cDNA derived from untreated *A.thaliana*, while the *MTA* genomic locus (*MTAgen*) was amplified from *A. thaliana* genomic DNA, both using primers designed for Gateway recombination. TuMV-GFP *3’Peak* was isolated from *p35S::TuMV-GFP*, similarly m1-m5 (small and big) were obtained by amplifying selected areas of *p35S::TuMV-GFP*. The control was obtained by amplifying overlapping primers

The PCR products were then cloned into *pD207* using Gateway™ BP Clonase™ Enzyme Mix (11789013, Invitrogen™) according to the manufacturer’s recommendations. Cloned products were introduced into One Shot™ TOP10 Chemically Competent *E. coli* (C404010, Invitrogen) through heat-shock transformation. Transformed colonies were selected via antibiotic resistance and confirmed through Sanger sequencing. Plasmids were purified using the QIAprep Spin Miniprep Kit (27104, QIAGEN) following the manufacturer’s protocol.

Finally, *pD207::mSeries*, *pD207::3’Peak*, and *pD207::MTAgen* were recombined into *pUBNGFP*, *pGdL5/pGdL3*, and *pGWB440*, respectively, using Gateway™ LR Clonase™ II Enzyme Mix (11791020, Invitrogen), generating: *pUBN::m#*, *pGdL5::3’Peak* and *pGdL3::3’Peak* and *pGWB440::MTAgen*. The introduction of the MTA locus in to *pGWB440* generated a 3’ YFP tagged version of *MTA* (hereafter *pMTA::MTA-YFP*). These constructs were subsequently transformed into *A.tumefaciens* GV3101 for further experiments.

### Plant material

Seedlings were all grown with a 16-h photoperiod and 8-h dark conditions at 23 °C (light) 22°C (dark) for both germination and growth. Arabidopsis lines *vir1*, *hakai* and *fip37* (Salk_138482, Salk_148797, Salk_018636, respectively) were obtained from the Brian Gregory lab at the University of Pennsylvania, USA. Reader’s mutants *ect2-1* (SALK_002225), *ect3-1*(SALK_077502), *ect4-2* (GK-241H02) (*ect4*)*, ect5*(SALK_111736c), *ect7*(SALK_105881c), *ect9-1* (SALK_030033c) (*ect9*) and the double and triple mutants *ect23, ect24* and *ect234* were obtained from Dr. Monika Chodasiewicz lab. *ect4-1* (SALK_151516), *ect4-3* (SALK_112012), *ect9-2* (SALK_030033), *ect9-3* (SALK_052750) were purchased from ABRC^132^.

*Col0 ABI3::Col0* and *mta ABI3::MTA* were obtained from Prof. Rupert Fray lab.^71^ The *MTA-YFP*-complement line was generated by introducing *pMTA::MTA-YFP* into *mta ABI3::MTA* mutant by Agrobacterium*-*mediated transformation through floral dipping.

*Nicotiana benthamiana* (Nicotiana) seeds were obtained from publicly available collections.

All seeds were sterilized before each experiment by shaking them in 70% EtOH with 0.05 percent SDS for 10 minutes, followed by 2 absolute EtOH washes. Seeds were left to dry on sterile filter paper in a laminar flood hood. Filter papers were then placed in a sealed petri dish to avoid contamination. Petri dishes containing the seeds were then incubated in the dark for 48 hours at 4 °C, and afterwards, seeds were sprinkled into presterilized, hydrated Jiffy pods (32170142, Jiffy®), agar plates or soil.

Jiffy pods were prepared the day before by hydrating the pods with 2 g/l of Folicat Plantifol 20-20-20 (Atlantica) in milliq-water overnight, followed by autoclave sterilization. Jiffy pods were then arranged in trails containing a total of 32 pods. Sown trails were then put in growing incubators PGC FLEX (140237, Conviron), covered with a lid for 7 days, and then seedlings were selected so only one seedling was present in each pod. Jiffy trails were then irrigated with tap water.

Plates were prepared by pouring autoclaved ½ MS media with 9 g/l of agar (A1296,SIGMA) in to square, 12 cm by 12 cm plates. ½ MS media was prepared by mixing 0.5 g/l of MES monohydrate(E169, VWR) and 2.2 g/l of Murashige and Skoog Basal Salt Mixture (M5524, Sigma); pH was then adjusted to 5.7.When necessary, selection markers were added in the autoclaved media before pouring.

Nicotiana seeds were germinated directly in pre-hydrated soil (potting soil A 240, Stendler) in lid-covered trails in a growth room at 22°C for 10 days. Seedlings were then selected so only one seedling would remain in each pot and irrigated with tap water.

### Virus propagation and infection procedures

Primary infections were induced by needleless agroinfiltration of 20-day-old Nicotiana plants with 500 µl of resuspended Agrobacterium in 10 mM MgCl, 10 mM Murashige and Skoog Basal Salt Mixture (M5524, Sigma) with a final O.D. of 0.6. The apical meristems of infected plants were then harvested at 10 dpi for subsequential infections. Plant tissues were ground at 4°C and resuspended in 2 volumes of pre chilled 0.1 M sodium phosphate buffer, pH 7.5.

Virus suspension was clarified first by Miracloth filtering (475855, Merck Millipore) and then by a 45 µm nylon syringe filter (15131499, Fisher Brand).

Viruses were quantified by GFP fluorescence using a 96-well plate on an infinite M200PRO (Tecan). The virus concentration was then adjusted and either used for analysis or to infect secondary hosts. Secondary infections were carried out by rubbing host leaves with 10 µl of viral resuspended solution using a flat-tip painters brush and silicon carbide powder. Infected plants were then harvested at different time points, frozen in liquid nitrogen, and stored at -80 °C.

### Plant sample preparation

Frozen plant tissue was ground using a mortar and pestle, frequently cooled by liquid nitrogen.

### DNA and RNA extraction

Frozen plant powder and viral suspensions were used for DNA and RNA extractions.

Genomic DNA was extracted using the DNeasy Plant Pro Kit (69204, Qiagen) following the producer’s recommendations.

Total RNA was extracted using Trizol Reagent (15596018, Invitrogen) following producers’ recommendations with some variations, briefly: 150 mg of frozen plant tissues or 150 µl of viral suspension were incubated for 3 min at room temperature with 1.5 ml ice-cold Trizol. Samples were then mixed with 0.3 ml of chloroform (288306, Sigma Aldrich) and incubated on a shaker for 3 minutes at room temperature. Samples were then centrifuged at 12500 g for 20 minutes at 4°C. The 0.6 ml of the aqueous translucid supernatant was retrieved and mixed with 0.17 ml of 3M sodium acetate (1.06264, Sigma Aldrich) and 0.77 ml of isopropanol (278475, Sigma Aldrich), incubated on a shaker for 5 min at room temperature, and left O/N at 4°C. Samples were then centrifuged at 12500 g for 15 minutes at 4°C, the supernatant was removed, and the RNA was resuspended in 0.75 ml of DEPC water (4387937, Thermo Fisher). Samples were then incubated on a shaker at RT for 5 min and then mixed with 0.75 ml of isopropanol and chilled on ice for 20 min. Samples were then centrifuged again at 14000 g for 20 min at 4°C, and then supernatant was removed. The RNA pellet was then washed twice with 75% EtOH and resuspended in DEPC water. Nucleic acid concentration was estimated via NanoDrop™.

Viral enriched RNA was purified using Plant Virus RNA Kit (PVR100, Geneaid) following the producers’ recommendations.

mRNA was purified using Dynabeads™ Oligo(dt)_25_ (61005, Thermo Fisher) following producers’ recommendations.

### m^6^A dot blotting

mRNA and virus enriched RNA were prepared in several sequential dilutions. 3 µl droplets were pipetted on a nylon membrane and let dry at room temperature for 5 to 10 min. RNA was then crosslinked using an Fisherbrand™ UV Crosslinker with Adjustable Height (13-245-222, Fisher Scientific) for 30 seconds twice, rotating the membrane to ensure uniform crosslinking. Membrane was then incubated for 2 h at room temperature in blocking solution: 5 percent Blotting-Grade Blocker (1706404, BioRad) in PBS-T. PBS-T was prepared by adding 0.02 % (v/v) Tween®20 (P9416, Sigma Aldrich) in PBS (524650, Sigma Aldrich). Then the membrane was let to incubate overnight in a horizontal shaker at 4°C with N^6^-Methyladenosine (m^6^A) Recombinant Rabbit Monoclonal Antibody (RM362, Thermo Fisher) diluted 1:5000 in blocking solution. The membrane was then washed every 10 minutes for 5 times with PBS-T and then incubated with 1:15000 Α-Rabbit IgG (H+L), HRP Conjugate (W4011, Promega) in blocking solution for 1.5 h at room temperature in a horizontal shaker. The membrane was then washed with PBS-T every 10 minutes 5 times and then developed using SuperSignal™ West Femto Maximum Sensitivity Substrate (34095, Thermo Fisher) according to manufacturer recommendations. Images were captured using ChemiDoc XRS+ System (BioRad).

### SDS page and Western blotting

Total sample proteins were obtained by boiling frozen plant tissue with a pre-stained 2X SDS loading buffer (100 mM Tris-Cl pH 6.8, 200 mM DTT, 4% (w/v) SDS, 0.2% (w/v) Bromophenol blue, 20% (v/v) Glycerol) at 85°C for 10 minutes. SDS page was carried out using Mini-PROTEAN® TGX™ Precast Gels (4561033, BioRad), run at 125V and 3.5 A for 1 hour. SDS page gels were washed twice with milli-Q water and then incubated O/N in SimplyBlue™SafeStain (LC6065, Life Technologies). SDS page gels subjected to Western blotting were first blotted using Trans-Blot Turbo Mini 0.2 µm PVDF Transfer Packs (1704156, BioRad) and then the membranes were blocked in a 3-5% blocking solution (Blotting-Grade Blocker,1706404, BioRad) in TBS-T buffer (10x Tris Buffered Saline, 1706435, BioRad; TWEEN^®^ 20, P9416, Sigma Aldrich) on an horizontal shaker for 2 hours at room temperature, followed by incubation with the designed primary antibody in blocking buffer on an horizontal shaker overnight at 4°C. The primary antibodies used in the study are as follow: α-GFP (ab290, Abcam), α-Tubulin (ab4074, Abcam), α-H3 (ab12079, Abcam), α-TuMV (Agdia). Membranes were then washed 5 times with TBS-T buffer every 10 minutes, and then incubated with a secondary antibody (W4011, Promega) for 90 minutes. The membrane was then washed 4 times with TBS-T buffer and developed using either Clarity Max Western ECL Substrate (BioRad, 1705062) or SuperSignal™ West Femto Maximum Sensitivity Substrate (34094, Thermo Fisher Scientific). For both Western blots and SDS page gels, images were recorded using the ChemiDoc XRS+ System (BioRad).

### m^6^A LC/MS analysis

LC/MS analysis was carried out following L.Mathur *et al.* protocol^89^, with some modifications:polyadenylated RNA or viral enriched RNA was first fragmented using Nuclease P1 from *Penicillium citrinum* (N8630, Sigma) and dephosphorylated with Phosphatase, Alkaline from *Escherichia coli* (P4069, Sigma) following the L.Mathur *et al.* protocol^89^. Then nucleosides were precipitated by centrifuging at 15000 g at 4°C for an hour. 40 µl of supernatant containing the RNA nuclesides were mixed with 80 µl of acetonitrile (271004, Sigma Aldrich). The sample order was randomized and then loaded into a UHPLC-IDEX Tribrid Orbitrap-Mass Spectometer (Thermo Fisher). Adenosine and m^6^A peaks were then analyzed using XCALIBUR™SOFTWARE. Species quantification was done by calculating the integral of the area below each peak. Calibration curves were done as specified by L.Mathur *et al.* protocol^89^, changing the percentage of acetonitrile from 25% to 33.33%.

### Protoplast purification and RNA stability assays

Protoplasts were isolated from 4-week-old Arabidopsis plants. Leaves 3 through 8 (six per plant) were harvested and cut into 0.5–1 mm strips. The cut strips were processed following the protocol by *Yoo et al.*^133^, with some modifications. Briefly, strips were submerged in 8 mL of Enzyme Solution (0.4 M mannitol, 20 mM KCl, 20 mM MES pH 5.7, 1.5% Cellulase R10, 0.4% Macerozyme R10, 0.05% BSA, and 10 mM CaCl_2_) in a 5 cm Petri dish, ensuring the strips remained submerged and non-overlapping. The tissue was vacuum-infiltrated for 10 min, incubated in the dark at room temperature (RT) for 2 h, and then placed on a horizontal shaker at 70 rpm for 20 min to release the protoplasts. The resulting suspension was filtered through a 100 µm mesh and centrifuged at 200g for 1 min at 4°C. The pellet was delicately resuspended in 2 mL of W5 buffer (154 mM NaCl, 125 mM CaCl_2_, 5 mM KCl, and 2 mM MES, pH 5.7). To ensure culture homogeneity, the suspension was pooled and aliquoted into round-bottom tubes. The protoplasts were then allowed to precipitate by gravity on ice for 40 min. Following two subsequent washes with W5 buffer via gravity precipitation, the cells were finally resuspended in MMg buffer (0.4 M mannitol, 15 mM MgCl_2_, and 4 mM MES, pH 5.7) to a final volume of 5.5 mL. For transformation, 10 µg of plasmid DNA was added to 10 mL round-bottom tubes, followed by 750 µl of the protoplast suspension. Negative controls received an equal volume of 10 mM Tris-HCl (pH 8.0). To each tube, 825 µl of PEG solution (40% w/v PEG 4000, 0.2 M mannitol, and 100 mM CaCl_2_) was added and mixed gently. Following an 8 min incubation at RT, the reaction was quenched by the slow addition of 3.3 ml of W5 buffer. The mixture was centrifuged at 200g for 2 min at 4°C, and the supernatant was completely removed to eliminate residual PEG toxicity. For each vector combination, the protoplasts preparations were resuspended in 800 µl of W1 buffer (0.5 M mannitol, 20 mM KCl, and 4 mM MES, pH 5.7) and 180 µl were aliquoted into 4 wells of a black clear bottom 96 well plate. The following day, for each vector combination, 20 µl of W1 buffer containing 0.5 M actinomycin D and 0.1 M α-amanitin was added to all wells except for one row (12 wells), which served as the control and received 20 µl of W1 buffer only. For each time point and vector combination, 12 wells were harvested for downstream analysis.

### Nuclear and cell fractionation

Subcellular fractionation and nuclei isolation were performed according to amodified protocol by *Gendrel et al.*^134^. Approximately 2 g of plant tissue was ground to a fine powder in liquid nitrogen and homogenized in 25 mL of Extraction Buffer 1 (0.4 M sucrose, 10 mM Tris-HCl pH 8.0, 10 mM MgCl_2_, 5 mM β-mercaptoethanol, and protease inhibitors). The homogenate was filtered through a 100µm nylon mesh into a pre-cooled 50 mL Falcon tube. The filtrate was centrifuged at 1,500g for 15 minutes at 4°C. Following centrifugation, the supernatant was collected and stored at –80°C as the cytosolic fraction, while the pellet was gently resuspended in 20 mL of Extraction Buffer 2 (0.25 M sucrose, 10 mM Tris-HCl pH 8.0, 10 mM MgCl_2_, 1% Triton X-100, 5 mM β-mercaptoethanol, and protease inhibitors) using a wide-bore tip. The suspension was incubated on ice for 10 minutes and centrifuged again at 1,500g for 10 minutes at 4°C. The resulting pellet was resuspended in 500 µl of Extraction Buffer 3 (1.7 M sucrose, 10 mM Tris-HCl pH 8.0, 2 mM MgCl_2_, 0.15% Triton X-100, 5 mM β-mercaptoethanol, and protease inhibitors) and carefully layered over an equal volume of Extraction Buffer 3 to create a sucrose cushion. After centrifugation at 16,000g for 1 hour at 4°C, the supernatant was discarded, and the nuclear pellet was resuspended in Nuclei Lysis Buffer (50 mM Tris-HCl pH 8.0, 10 mM EDTA, 0.1% SDS, and protease inhibitors). Samples were either used immediately or flash-frozen in liquid nitrogen for downstream applications.

### Real-time qPCR

cDNA was synthesized from total RNA using SuperScript™ III First-Strand Synthesis SuperMix (18080400, Thermo Fisher) following the random hexamer protocol. RT-qPCR was performed using SsoAdvanced™ Universal SYBR^®^Green Supermix (1725271, BioRad), following the supplier specifications for designing the thermocycle. Samples were run on a CFX384 Touch Real-Time PCR Detection System (BioRad). Uniformity between the samples was achieved by RNA quantification and calibration before cDNA synthesis. At least 3 biological replicates and 2 technical replicates were included for each run. Fold changes were calculated using the 2^-ΔΔCt^ method. Arabidopsis thaliana AT5G15710 and AT1G13320 were used as reference genes, the uninfected plants for each line were used as reference sample. TuMV VPg was used as a target gene to evaluate viral titers. Primer sequences are listed in Supplementary Table 1.

### Me-RIP sequencing

m^6^A IP and sequencing were done by Novogene. Briefly, poly-adenylated RNA was purified from total RNA, derived from 12 dpi TuMV infected *WT* and *MTA-YFP* samples. Each condition was provided with 3 biological replicates, each derived from the RNA collected from 10 different plants. Poly-adenylated RNA was then fragmented in to smaller transcripts, then while a fraction was kept as input, the rest was immunoprecipitated with m^6^A specific antibody (done by Novogene). The IP and input fragments were then added adaptors and used for next gen sequencing using an illumina novaseq x plus instrument (https://www.illumina.com/) with paired-end.

### Transcriptome analysis

Arabidopsis *Col-0/ABI3::Col0* (WT) and *mta/ABI3::mta* (*mta*) plants, either mock-treated (WT and *mta*) or infected with TuMV-GFP (WT), were harvested at 12 dpi for RNA-seq. For each condition 3 biological replicates were provided each constituted by total RNA collected form 10 plants. Transcriptome sequencing was performed by Novogene using an Illumina NovaSeq X Plus instrument (https://www.illumina.com/) with paired-end.

### Sequencing data analysis

Raw reads from transcriptome and Me-RIP sequencing underwent quality control and adapter removal using FastQC^135^ (https://www.bioinformatics.babraham.ac.uk/projects/fastqc/) and TrimGalore^136^ (https://www.bioinformatics.babraham.ac.uk/projects/trim_galore/), respectively. Clean transcriptome reads were mapped to the *Arabidopsis thaliana* genome (GCA_000001735.1) using TopHat2^137^. Transcriptome differential gene expression and FPKM values were calculated using Cufflinks^138^ (https://cole-trapnell-lab.github.io/cufflinks/) and DESeq2^139^. For Me-RIP, paired-end reads were mapped using Bowtie2^140,141^ (https://bowtie-bio.sourceforge.net/bowtie2/manual.shtml). Me-RIP peak calling was performed using MACS2^142^ (https://genomebiology.biomedcentral.com/articles/10.1186/gb-2008-9-9-r137), and annotated by ChIPseeker R package (v1.36.0)^143^, which maps modification sites to transcript features (5′ UTR, CDS, 3′ UTR, etc). Biological replicates were then merged using Multiple Sample Peak Calling Tool (MSPC, https://genometric.github.io/MSPC/)^144^, using the following pipeline: *mspc -i Rep1_peaks.bed Rep2_peaks.bed Rep3_peaks.bed -r Bio - w 1e-4 -s 1e-8 -o result_macs_peaks* Final data processing was performed using deepTools^145^ (https://deeptools.readthedocs.io/en/develop/).

### ONT direct RNA sequencing and data analysis

#### Sample Preparation and Sequencin

vRNA was extracted from *A.talliana* WT and *mta* mutant plants, infected with TuMV-GFP at 3 dpi. For each condition we analysed 3 biological replicates, with each resulting from RNA collected from 10 plants. Direct RNA sequencing was performed using Oxford Nanopore Technology (ONT) on a PromethION 24 platform, with a PromethION flow cell for RNA (FLO-PRO004RA). Sequencing libraries were prepared following the manufacturer’s protocol for direct RNA sequencing (ONT, Oxford, UK). Raw FASTQ files were generated from the basecalled reads.

#### Read Mapping

For mapping of ONT reads, we used the following approach:

#### Mapping to TuMV genome

ONT reads were aligned to the TuMV-GFP coding sequence (CDS) reference (*TuMV-GFP_cds.fa*) using minimap2 (v2.26)^146^ with the -ax map-ont preset, which is optimized for ONT reads. The command used was: *minimap2 - t 30 -ax map-ont /path/to/TuMV-GFP_cds.fa /path/to/combined.fastq.gz > aligned.sam* **Mapping to *Arabidopsis* genome:** For host transcriptome analysis, ONT reads were additionally mapped to the *Arabidopsis thaliana* reference genome (GCA_000001735.1) u sing the same minimap2 parameters. Only uniquely mapped reads were retained for downstream analysis.

#### m⁶A Modification Calling

Methylated adenine sites were identified using m6Anet (v2.1.0)^147^, a machine-learning-based tool that predicts m⁶A modifications from ONT direct RNA sequencing data. m6Anet calculates a probability of modification for each adenine position based on the raw electrical signal and the surrounding nucleotide sequence (k-mer context).

The output file (data.site_proba.csv) contained the following columns for each predicted site:

- transcript_id – transcript identifier
- transcript_position – position within the transcript
- n_reads – read coverage at that position
- probability_modified – posterior probability of m⁶A modification
- kmer – 5-mer sequence centered on the adenine
- mod_ratio – estimated proportion of modified reads

#### Training of m6Anet on Viral Data

Because m6Anet’s default model is trained on cellular transcriptomes, its accuracy on viral RNA sequences can be limited. To improve discrimination of true m⁶A sites from false positives in the TuMV genome, we retrained the m6Anet model using a custom training dataset derived from ONT reads of viral origin. This viral-trained model accounted for the distinct nucleotide composition and secondary structure of TuMV RNA, resulting in more reliable m⁶A predictions.

#### Integration with Me-RIP Data

The normalized ONT m⁶A probabilities were then plotted alongside the Me-RIP peak signals using deepTools^145^ and custom R scripts (ggplot2)^148–150^. This integrated approach allowed us to compare single-nucleotide resolution m⁶A calls (ONT) with enrichment-based peaks (Me-RIP).

#### Motif discovery

The HOMER suite’s *findMotifsGenome.pl* tool (http://homer.ucsd.edu/homer/motif/) was used for the *de novo* motif discovery. Motifs with a p-value < 10^-2^ were considered significant. Significant motifs were then mapped onto the TuMV genome using HMMER (http://hmmer.org/) to identify potential methylation sites within the TuMV genome.

#### Heatmaps and gene ontology analysis

Heatmaps and clusters were generated using DESeq2^139^ (https://bioconductor.org/packages/release/bioc/html/DESeq2.html) and pheatmap (https://www.rdocumentation.org/packages/pheatmap/versions/1.0.12/topics/pheatmap)^151^. Gene ontology analysis was performed using AgriGO (https://systemsbiology.cau.edu.cn/agriGOv2/)^152^. All plotting and secondary data analysis were performed using RStudio^148–150^.

#### Data analysis and plotting

Data analysis was performed using RStudio^148,153^, and all figures were generated with the ggplot2 package^149,150^. The analysis followed standard workflows and functions available in the cited packages; custom scripts are available from the corresponding author upon reasonable request.

## RESULTS

### TuMV-GFP vRNA titer and protein expression are regulated by host m^6^A methylation

To study the role of m^6^A in plant viruses, we used a GFP-tagged TuMV cDNA cloned into an *Agrobacterium*-compatible plant expression vector (*p35S::TuMV-GFP*). The GFP-tagged TuMV ensures consistent results and facilitates monitoring of the infection process^69^ (Fig 1A). GFP protein levels were used to estimate viral protein synthesis since GFP behaves as one of the viral cistrons. The virus was multiplied in *Nicotiana* plants through agrobacterial transient expression before infecting the subsequent hosts like Arabidopsis.

**Figure 1:**
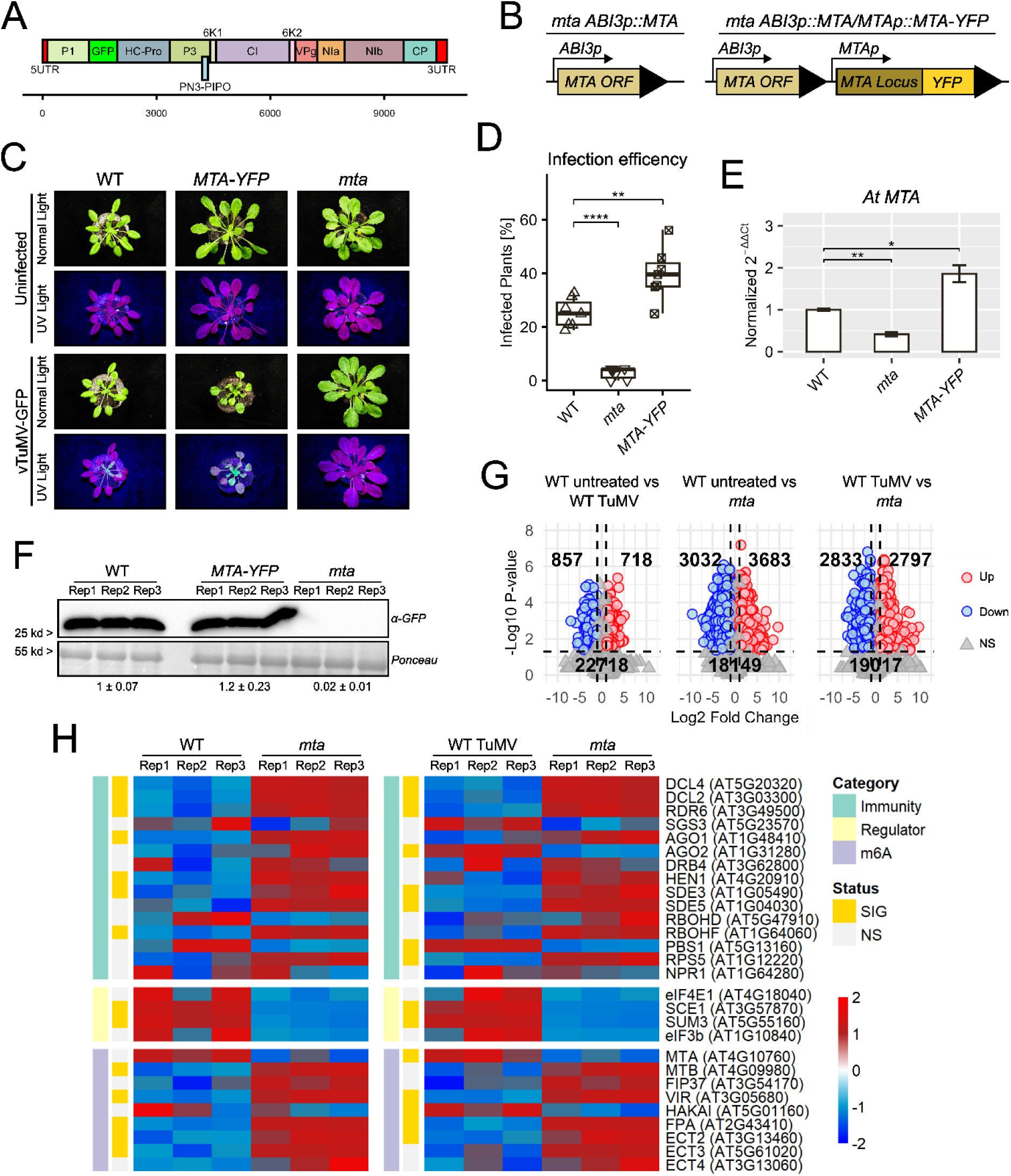
***mta* mutant plants show resistance to TuMV infections** (A) Schematic representation of the *TuMV* genome structure, highlighting the insertion of the GFP reporter. (B) Schematic representations of the expression cassette in the *mta ABI3::MTA* mutant and in *MTA-YFP* complemented line. (C) Panel showing WT, *MTA-YFP* and *mta*, infected with TuMV-GFP at 12dpi. (D) Infection frequency in vTuMV-GFP-infected samples. Plants were classified as infected if they exhibited GFP fluorescence between 12 and 20 dpi. Percentages were calculated as the ratio of symptomatic plants to the total number of infected plants. Statistical comparisons between specified groups were performed using two tailed Student’s t-tests. Significance levels are indicated by asterisks (“*”), where the number of asterisks corresponds to -log_10_(p-value), with a minimum threshold set at p-value < 0.05. Non-significant comparisons are not labelled. (E) RT-qPCR analysis of *At MTA* expression in WT, *MTA-YFP* and *mta*, relative to WT. Results are shown as normalized 2^-ΔΔCt^. Each value represents the mean of six biological replicates, with three technical replicates per biological sample. Error bars represent the variance of each group. Statistical comparisons between specified groups were performed using two tailed Student’s t-tests. Significance levels are indicated by asterisks (“*”), where the number of asterisks corresponds to -log_10_(p-value), with a minimum threshold set at p-value < 0.05. Non-significant comparisons are not labeled. (F) Western blot analysis of TuMV-GFP-infected samples using α-GFP antibodies at 12 dpi. Band intensities were quantified relative to the most intense WT band, with values representing fold changes. Each band corresponds to a biological replicate, with each replicate consisting of four plants. (G) Volcano plots showing upregulated, downregulated, and non-significant genes from comparisons between untreated WT, infected WT, and *mta*. (H) Heatmap representing the expression profiles of select antiviral, regulatory, and m^6^A-related genes across WT and *mta* mutant backgrounds. Expression levels are depicted as row-wise Z-scores calculated from log_2_-transformed counts; values are capped between –2 and 2 to highlight relative expression differences. Statistical significance was determined by a two-tailed Student’s t-test for each gene, followed by Benjamini-Hochberg adjustment for multiple testing. Genes with an adjusted p-value < 0.05 are indicated in the status annotation (yellow).

Infected tissues exhibiting visible GFP fluorescence were harvested at different time points (Fig. S1A-D). GFP was primarily detected as a 27 kDa protein (Fig. S1B), and subsequent viral purification coupled with qPCR analysis of key viral genes like *VPg* confirmed that fluorescence levels correlated with viral titer in wild-type plants (Fig. S1C). GFP fluorescence became visually detectable at 12 days post-infection (dpi) in *Arabidopsis* ecotype Col-0; therefore, this time point was used for screening TuMV-GFP infections.

The homozygous mutants of the primary m^6^A methyltransferase of the writer complex, *MTA* is lethal, making *mta* mutants challenging. We used the viable *mta* mutant line (*mta ABI3p::MTA*, hereafter *mta*), where *MTA* expression is restricted to embryonic development by the *ABI3* promoter. As a wild-type control, we used *Col-0 ABI3p::Col-0* (hereafter WT), which was transformed with an empty cassette containing only the *ABI3* promoter^70^.

We also generated a complementation line in the *mta* background (*MTAp::MTA-YFP/mta*, hereafter MTA-YFP), in which the *MTA* locus was fused to a YFP reporter (Fig 1B). While the WT phenotype was completely restored in MTA-YFP, *MTA* expression levels appeared to be higher than in the WT, providing us with a mild overexpressor (Fig 1E).

Twenty-day-old WT, *mta* and MTA-YFP plants were infected with vTuMV-GFP harvested from infected *Nicotiana* plants, and fluorescence was monitored at 12 dpi. Only 2.7% of *mta* plants showed fluorescence, compared to 25.2% in WT and 39.8% in MTA-YFP (Fig. 1C,D). Consistently, GFP signal remained undetectable in *mta* indicating an almost complete absence of viral protein levels (Fig 1F).

Writer mutants like *mta* have been previously reported to exhibit primed-immunity and autoimmunity phenotypes^71–73^, which might interfere with viral infections outputs. To investigate this possibility, we performed transcriptome sequencing on *mta*, WT, and TuMV-GFP-infected WT at 12 dpi. Principal component analysis (PCA) revealed a total variance of 45.79% in PC1 and 19.51% in PC2, separating infected and uninfected WT samples, as well as *mta*, into distinct groups (Fig. S1E). A total of 857 genes were significantly downregulated and 718 genes upregulated between uninfected and infected WT. However, the most substantial differences in gene expression were observed when comparing either untreated or infected WT to *mta* (Fig. 1G).

We first attempted to identify these genes through Gene Ontology (GO) analysis. Although we found some categories related to immune defense, none of these genes have been reported to possess antiviral functions (Fig. S1G). We then chose to investigate key genes involved in the viral immune response, regulation of viral replication and proliferation, and members of the m^6^A writer and reader complexes (those not shown did not exhibit significant differential expression). As expected, *mta* showed upregulation of several genes involved in viral immunity and downregulation of various regulators, making *mta* a potential candidate for primed innate immunity. Conversely, m^6^A-related genes were mostly upregulated, likely as a compensatory effect for the loss of *MTA*.

As *mta* exhibits a pleotropic phenotype, we tested other mutants of the m^6^A writer machinery in *Arabidopsis* to confirm the m^6^A-specific effect on viral infection. We analyzed *vir1*, *hakai*, and *fip37* mutants, alongside *mta*, MTA-YFP, and WT, observing fluorescence at 12 dpi. Similar to our observations in *mta*, none of the writer mutants showed visible fluorescence at 12 dpi or even at 20 dpi (Fig 2A). To determine if a compromised writer complex was causing resistance or resilience to TuMV-GFP, we examined an earlier time point of 3dp. Interestingly, the viral RNA (vRNA) titer in these lines at 3 dpi was, albeit lower, not significantly different from WT in most cases, with the exception of *vir1* (Fig 2B).

**Figure 2:**
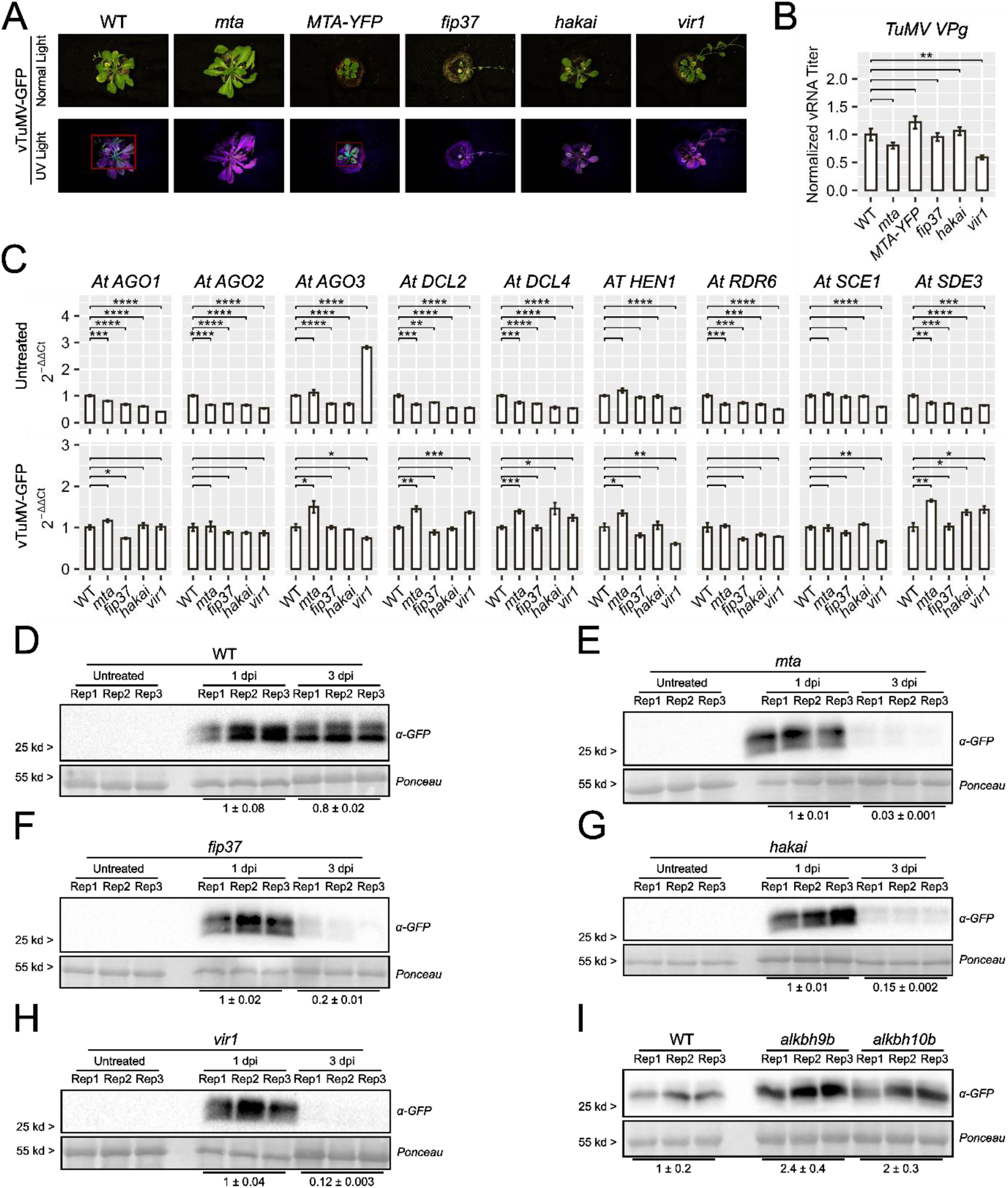
TuMV protein expression is modulated by the host writer complex. (A) Panel showing WT, *mta*, *MTA-YFP*, *fip37*, *hakai*, and *vir1*, infected with TuMV-GFP at 20 dpi. (B) RT-qPCR quantification of the viral titer, using *TuMV VPg* as target, normalized against WT, at 3 dpi. Each value represents the mean of six biological replicates, with three technical replicates per biological sample. Error bars represent the variance of each group. Statistical comparisons between specified groups were performed using two tailed Student’s t-tests. Significance levels are indicated by asterisks (“*”), where the number of asterisks corresponds to -log_10_(p-value), with a minimum threshold set at p-value < 0.05. Non-significant comparisons are not labeled. (C) RT-qPCR analysis of immunity genes expression in WT, *mta*, *fip37*, *hakai*, and *vir1*, relative to WT at 3 dpi. Results are shown as normalized 2^-ΔΔCt^. Each value represents the mean of six biological replicates, with three technical replicates per biological sample. Error bars represent the variance of each group. Statistical comparisons between specified groups were performed using two tailed Student’s t-tests. Significance levels are indicated by asterisks (“*”), where the number of asterisks corresponds to -log_10_(p-value), with a minimum threshold set at p-value < 0.05. Non-significant comparisons are not labeled. (D-H) Western blot analysis of TuMV-GFP-infected WT and m^6^A mutants using α-GFP antibodies at different time points. Band intensities were quantified relative to the most intense WT band, with values representing fold changes. Each band corresponds to a biological replicate, with each replicate consisting of four plants. (I) Western blot analysis of TuMV-GFP-infected WT and *alkbh* mutants using α-GFP antibodies at 3 dpi. Band intensities were quantified relative to the most intense WT band, with values representing fold changes. Each band corresponds to a biological replicate, with each replicate consisting of four plants.

One of the primary antiviral defenses in plants occurs through post-transcriptional gene silencing (PTGS), which was found to be upregulated in *mta* via transcriptome analysis (Fig 1H). We performed RT-qPCR for these PTGS-related genes using untreated and 3 dpi infected samples. With few exceptions, we observed lower expression in all mutants compared to WT in untreated samples. Furthermore, in infected samples, the expression of these genes was not consistently upregulated across all mutants, effectively ruling out a primed immune system as the primary resistance factor (Fig 2C).

To elucidate the molecular events underlying early infection, we monitored infection dynamics from 1 to 3 days post-infection. We observed a strong signal at 1 dpi in all mutants and the WT, but the signal was either absent or significantly faded at 3 dpi in all writer mutants (Fig 2D-2H). Additionally, all mutants displayed lower band intensities compared to WT at 1 dpi, indicating overall lower levels of viral protein synthesis (Fig. S2A).

Recently, *Sha et al.* demonstrated the recruitment of eraser *BjALKBH9B* by TuMV, which seems to exhibit an opposite behavior to what we observed in *Arabidopsis*^68^.

In order to clarify this phenomenon, we also tested *alkbh9b* and *alkbh10b* mutants, which both showed intense GFP signals at 3 dpi, revealing the opposite effect to what was previously reported in *B.juncea* (Fig 2I).

Since our experiments indicated either an enhanced vRNA stability or protein synthesis, we decided to investigate the m^6^A readers which play an important role in mediating these m^6^A- regulated processes^74–81^. We infected single (*ect2-1, ect5, ect7, ect9*), double (*ect23, ect24*), and triple (*ect234*) mutants with TuMV-GFP and collected samples at 2 dpi. Western blot revealed significant differences in GFP levels where *ect5, ect7,* and *ect9* mutants showed reduced band intensities, while *ect234, ect2 ect23* and *ect24* mutants displayed increased GFP levels relative to WT. These results suggest a dual role for ECTs, with ECT2, ECT3, ECT4 acting as negative regulators, while ECT5, ECT7 and ECT9 enhancing viral protein expression (Fig S2B).

We then challenged *ect2, ect4* and *ect9* with TuMV-GFP and monitored the visible GFP fluorescence at 12 dpi. Interestingly, all mutants including *ect9* showed similar levels of GFP fluorescence, signifying that while at early time points ECTs have a major role in determining viral protein production, none of these ECT mutants provideses plant virus resilience (Fig S2B).

We then decided to see if these mutants where affecting vRNA titers or if they affected viral proteins synthesis. For this we collected infected WT, *ect2, ect4* and *ect9* at 3 dpi. Similar to our previous results (Fig S2B), GFP levels in *ect2* and *ect4* were significantly higher than WT whereas *ect9* showed lesser GFP intensity (Fig S2D). Interestingly, vRNA levels showed substantial variations across the different mutants, with WT and *ect9* showing the same titer and *ect2* showing a 4-fold increase compared to WT. On the other hand, *ect4* had significantly reduced vRNA titers compared to WT.

Given the miscellaneous results of these ECT mutants as well as the poorly understood role that ECTs have in viral infections, we decided to include several allelic variants of *ect4* and *ect9* into the study and challenged them with TuMV-GFP. Samples were collected at 3 dpi and viral protein levels were measured by Western blotting. While band intensity varied significantly between the different alleles, the trend persisted having *ect4-1* and *ect4-3* showing higher whereas *ect9-2* and *ect9-3* lower band intensities.

Together, we conclude that TuMV infection in *Arabidopsis* is heavily influenced by the host m^6^A methylation. While m^6^A writers are necessary for viral protein synthesis, ECT readers differentially modulate vRNA titers and viral protein accumulation at early infection stages.

### TuMV is a direct target of m^6^A modification

While it was demonstrated that TuMV could be m^6^A modified^68^, further investigation was warranted given the differing results we obtained. To this end, we initially opted for MeRIP-sequencing^82,83^ using mRNA purified from TuMV-GFP infected WT and *MTA-YFP* plants at 12 dpi. Interestingly, while the number and position of the peaks remained consistent, we observed two significantly different peak profiles when comparing WT to *MTA-YFP*. *MTA-YFP* samples show a strong and consistent enrichment at the 3’-terminal region of the viral genome, characterized by a sharp, high-intensity peak across all replicates. In contrast, WT samples displayed a broader and more diffuse distribution of MeRIP signals with lower peak intensities and less pronounced enrichment in this region (Fig 3A). These peaks were primarily localized within viral CDS, overlapping with the regions corresponding to the *VPg*, *NIa*, *NIb*, and *CP* cistrons, as well as the 5′-most portion of the 3′UTR (Fig 3A). This distribution demonstrates that m^6^A modifications are predominantly concentrated toward the 3′ end of the TuMV genome.

**Figure 3:**
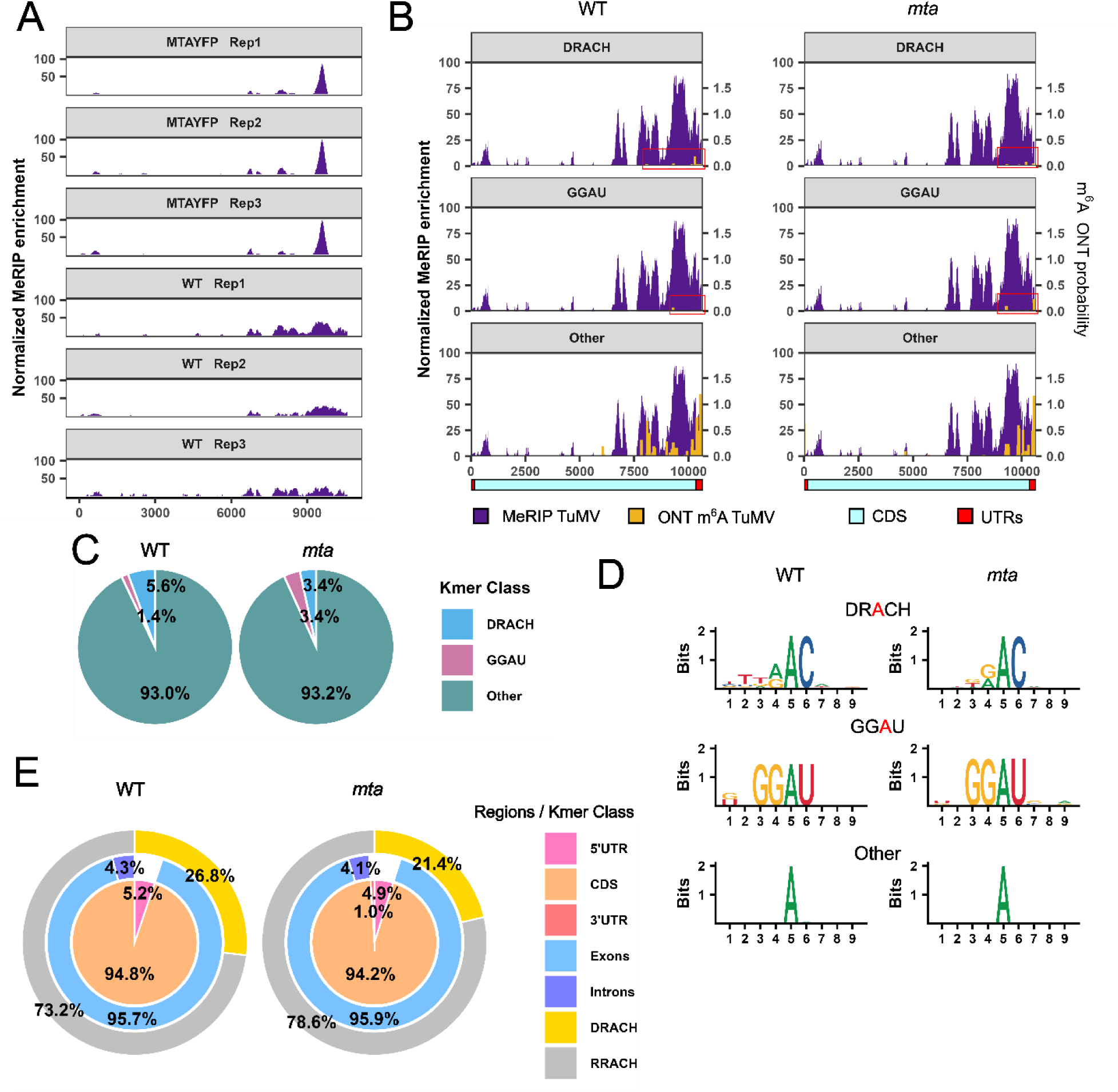
Distinct methylation patterns of TuMV vRNAs and host mRNAs. (A) MeRIP-sequencing peak profiles for TuMV-GFP infected WT and MTA-YFP at 12 dpi. Biological replicates are shown, with peak intensity normalized against the strongest signal. A total of three biological replicates per line were sequenced, each containing mRNA pooled from ten different plants. (B) Comparison between the average MeRIP-sequencing profile of WT and the average ONT direct RNA sequencing m^6^A profiles of TuMV-GFP infected WT and *mta* at 3 dpi. ONT peaks were generated by retaining only significant values and normalizing by read depth and the number of hits across biological replicates. A total of three biological replicates per line were sequenced, each containing vRNA pooled from ten different plants. The cartoon illustrates the genomic positions of each modification site relative to the TuMV-GFP genome. (C) Pie charts representing the relative ratios of the consensus k-mers observed via ONT sequencing of TuMV-GFP vRNA for each examined line. (D) Nucleotide logos representing the three k-mer consensus classes identified in the ONT analysis. Each logo was calculated independently for each plant line. (E) Pie charts representing the distribution of consensus k-mers across different RNA regions, as observed via ONT sequencing of host RNA impurities present in vRNA preparations for each examined line.

We attempted to use the HOMER motif finder to identify potential consensus sequences in these areas. However, most of the high-scoring motifs did not reflect canonical m^6^A consensuses, and some lacked significant adenosine residues (Fig S3).

Given that immunoprecipitation-based techniques can often yield false positives, we further investigated these sites using Oxford Nanopore Technologies (ONT) direct RNA sequencing^84,85^. For this experiment, we used TuMV-GFP infected WT and *mta* at the early time point of 3 dpi to ensure the presence of viral transcripts in the *mta* mutant. To avoid chip saturation with host transcripts, we utilized a Plant Virus RNA Kit, which yielded a five-fold increase in vRNA titers compared to traditional total RNA extraction methods (Fig S4A). To minimize false positives, m^6^A probability scores were normalized by read depth and the number of biological replicates supporting a modification at each site. We retained only those modifications present in at least two replicates, further filtering for significance by applying a threshold based on the global median value.

We identified m^6^A methylation in purified viruses from both WT and *mta* which may reflect residual methylation activity, as indicated by our previous dot blot experiments detecting trace methylation in mRNA from untreated *mta* (Fig S4B). Indeed, the *mta* mutant exhibited reduced overall methylation, characterized by a 20% reduction in the number of sites and a 57% decrease in the m^6^A probability score when compared to WT. We also observed a decrease in “DRACH/RRACH” (hereafter referred to as DRACH) and an increase in “GGAU” consensus sites. Unexpectedly, the vast majority of m^6^A modifications, and those with the highest probability scores, occurred at non-canonical sequences, labeled as “Others”; which are present in similar ratios in both WT and *mta* (Fig 3C). While the consensuses for DRACH and GGAU matched previous reports, the “Other” sites exhibited no discernible consensus (Fig 3D). To determine if these non-canonical sites were technical artifacts, we mapped the host transcripts present as impurities in our vRNA preparations. Most host modifications occurred within the CDS and 3’UTR of known genes, yet notably, no “GGAU” or “Other” motifs were detected in these cellular RNA fractions (Fig 3E). These results indicate that the non-canonical m^6^A modifications are virus-specific and suggest that TuMV transcripts are subject to a distinct methylation landscape compared to host mRNAs.

### m^6^A modification of TuMV occurs in the host nucleus

The writer complex is primarily nuclear-localized, and most host transcripts are immediately methylated post-synthesis^8,71^. However, positive-sense single-stranded RNA (+ssRNA) viruses do not require nuclear transcriptional activity to replicate, as most of their replication cycle is expected to occur in the cytoplasm^62,86–88^. Since our previous experiments indicated that the writer complex plays a major role in influencing TuMV-GFP protein expression, we decided to investigate the nuclear and cytoplasmic fractions of infected plants separately (Fig 4A). For this, we proceeded to separate nuclear and cytoplasmic fractions using TuMV-GFP infected WT plants at 2 and 3 dpi.

**Figure 4:**
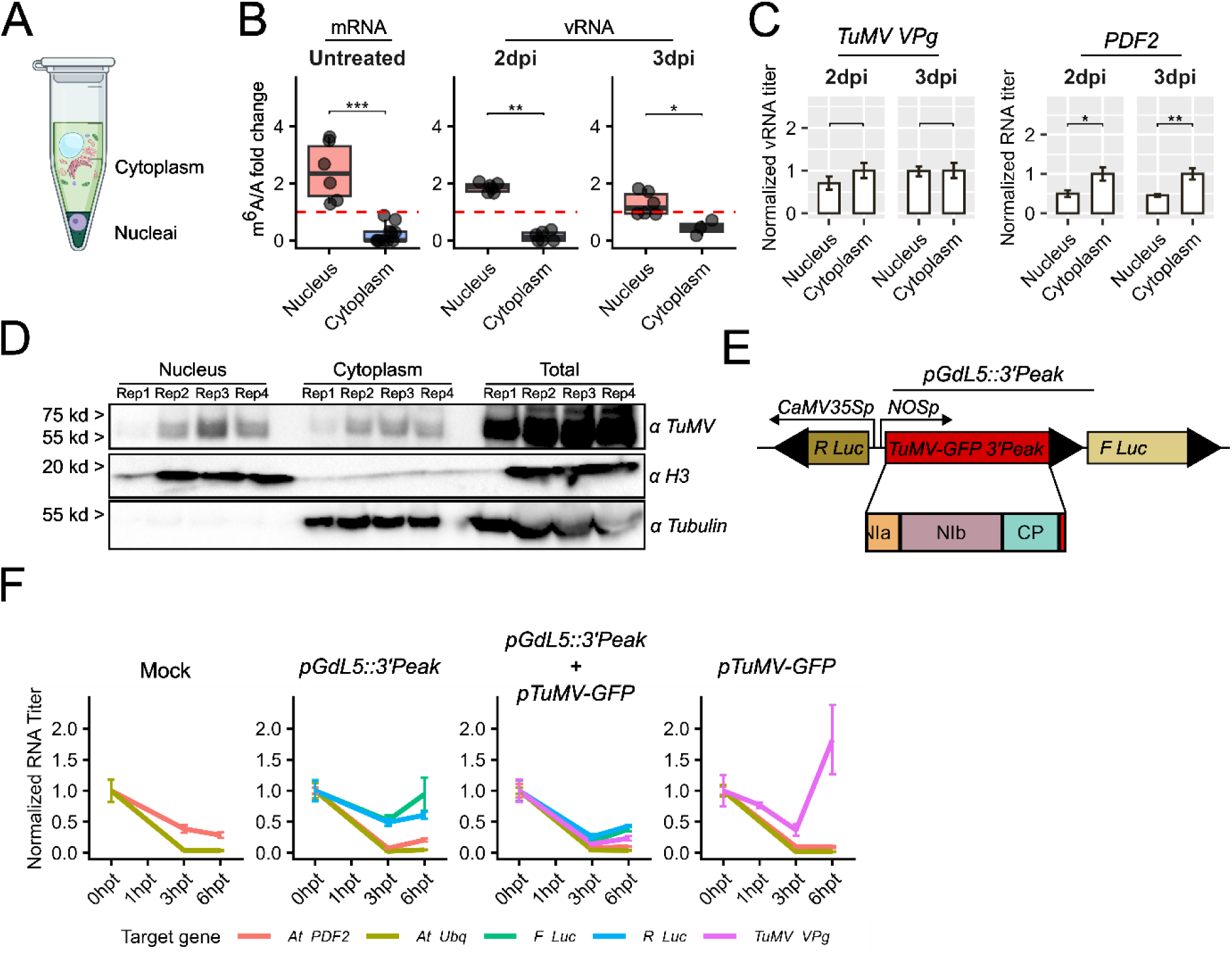
TuMV-GFP vRNAs are initially methylated in the nucleus. (A) Schematic representation of the nuclear/cytoplasm fractioning. (B) LC/MS quantification of m^6^A vs Adenosine (m^6^A/A). mRNA was purified from untreated WT samples while vRNA was obtained from infected plants using Plant Virus RNA Kit. (C) RT-qPCR quantification of the viral titer, using *TuMV VPg* as target, normalized against the cytoplasmatic fraction, at 2 and 3 dpi. Each value represents the mean of six biological replicates, with three technical replicates per biological sample. Error bars represent the variance of each group. Statistical comparisons between specified groups were performed using two tailed Student’s t-tests. Significance levels are indicated by asterisks (“*”), where the number of asterisks corresponds to -log_10_(p-value), with a minimum threshold set at p-value < 0.05. Non-significant comparisons are not labeled. (D) Western blot analysis of TuMV-GFP-infected WT at 3 dpi. We used α-TuMV antibodies to identify viral proteins, α-H3 for nuclear fractions and α-Tubulin for cytoplasmatic fractions. Band intensities were quantified relative to the most intense WT band, with values representing fold changes. Each band corresponds to a biological replicate, with each replicate consisting of four plants. (E) Schematic representation of the *pGdL5* and *pGdL3* expression system, showing both the reporter genes and the cassette position. (F) RT-qPCR analysis of the expression levels of *Fluc, Rluc, PDF2* and *TuMV VPg* in transformed protoplasts, over the course of the experiment, reported in hours post-treatment (hpt). Results are shown as normalized RNA titer, normalized against each target titer before the addition of actinomycin D and α-amanitin. Each value represents the mean of six biological replicates, with three technical replicates per biological sample.

To measure the methylation levels of these fractions, we utilized an LC/MS based method ^89^, using mRNA from untreated WT plants as a control. The m^6^A/A ratios clearly showed significantly higher m^6^A ratios in nuclear vRNAs than those of the cytosolic fraction at both 2 and 3 dpi, similar to that of host mRNA, (Fig 4B). Furthermore, the abundance of the viral *VPg* transcripts was assessed by qPCR in vRNAs isolated from host nuclear and cytosolic fractions (Fig 4C). TuMV-GFP vRNAs were clearly present at similar levels in the nuclei and cytoplasm of infected plants, while host specific transcripts like *PDF2* were predominantly cytoplasmic (Fig 4C). We also investigated whether any viral proteins localized to the host nucleus. Using a polyclonal antibody against TuMV, Western blot analysis of nuclear and cytoplasmic protein extracts revealed that TuMV proteins are present in both fractions, with a stronger signal observed in the nucleus than in the cytoplasm at 3dpi. (Fig 4D).

As a primary function of m^6^A is to regulate RNA stability^75,81,90,91^, we asked whether the reduced viral protein accumulation observed in writer mutants results from decreased vRNA stability. To test this, we employed a dual-reporter system containing the Firefly luciferase (Fluc) and Renilla luciferase (Rluc)^92^. The 3′ m^6^A hotspot region of 2729 bp of the vRNA (ranging from position 7800 - 10528, stop codon was removed) was fused either upstream (pGdL5) or downstream (pGdL3) of the FLuc reporter, and expressed in WT or *mta* protoplasts (Fig. 4E).

WT-derived protoplasts were transformed with *pGdL5::3’Peak*, *pTuMV-GFP*, or a combination of both, as the 3’Peak area encompasses parts of *NIa*, *NIb* and *CP*, potentially exerting RNA-stabilizing or partial catalytic effects. After overnight transformation, protoplasts were treated with 50 µM actinomycin D and 10 µM α-amanitin. Samples were collected at multiple time points, indicated as hours post-treatment (hpt). Host transcript levels like *PDF2*, *Ubq* decreased rapidly post-treatment in all cases; however, no significant difference was observed between *Rluc* and *Fluc* stability, albeit their titer decreased overtime (Fig 4F). Interestingly, the co-transformation of *pGdL5::3’Peak* and *pTuMV-GFP* exerted an overall negative effect on RNA stability. In contrast, protoplasts transformed with *pTuMV-GFP* alone exhibited initial decrease in vRNA titers, followed by a burst at 6h. While the latter trend is expected, given that the TuMV NIb replicase functions independently of the host transcriptional machinery, which are inhibited by actinomycin D and α-amanitin, we initially observed a transient decrease in TuMV-GFP vRNA titers. This early drop likely reflects the immediate impact of the host’s baseline antiviral defense. However, as these host pathways are subsequently suppressed or fail to keep pace with the infection, the viral titers exhibit a steady increase.

In contrast, the precipitous drop in RNA titers observed in the co-transformed samples likely results from vector overload. This intense competition for host resources, coupled with the inherent lytic activity of TuMV, likely destabilizes the cellular environment and leads to a loss of transcript integrity despite the breakdown of the host defense system (Fig 4F).

Together, we demonstrated the presence of TuMV-GFP vRNA and viral proteins within the nuclear fractions of the host cell, where they are positioned to interact with or to be targeted by the m^6^A writer complex. Furthermore, the specific methylation regions identified in TuMV-GFP do not significantly enhance transcript stability. This suggests that the primary role of m^6^A in the TuMV life cycle may be centered on the regulation of viral protein synthesis or translation efficiency rather than the protection of viral RNA from degradation. These findings reveal a previously unrecognized nuclear phase of the TuMV infection cycle and highlight a specialized role for the host epitranscriptomic machinery in modulating potyviral gene expression.

### Structural context and multi-site clusters define the TuMV methylation landscape

While nuclear localization and high m^6^A levels in the nucleus might explain canonical “DRACH” motifs, the nature of the “Other” sites remained unclear. To characterize these, we applied hierarchical clustering based on sequence similarity, identifying three distinct clusters independently for each line (Fig 5A). Cluster 1, comprising the majority of k-mers in both lines, showed no clear consensus other than the central adenosine target. Notably, clusters 2 and 3 displayed distinct nucleotide logos when comparing WT to *mta* (Fig 5A). Cluster 1 was more prominent in *mta* (82%) than in WT (71%), suggesting a higher degree of stochasticity in methylation placement in the mutant. The significant divergence between WT and *mta* logos for clusters 2 and 3 suggests they are likely deposited by distinct enzymes or that they are placed within different environments (Fig 5B-C).

**Figure 5:**
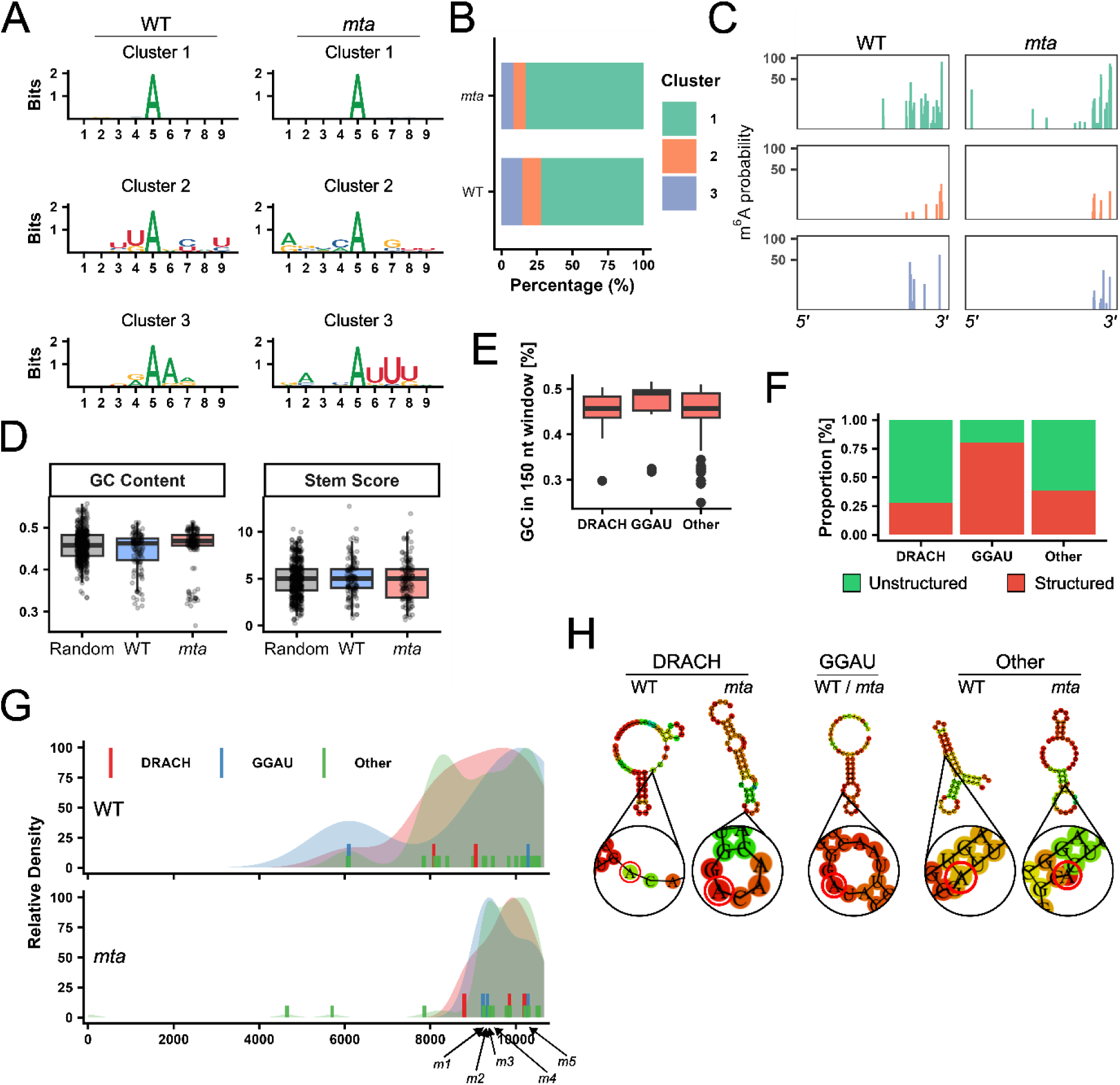
TuMV “Other”m^6^A methylation is influenced by the proximity to DRACH/GGAU sites and RNA secondary structure. (A) Nucleotide logos representing the three k-mer consensus clusters identified in the “Other” consensuses analysis. Each logo was calculated independently for each plant line. (B) Bar plot showing the different percentages of “Other” clusters in each line. (C) Spatial distribution of “Other” consensuses clusters in each line. (D) GC percentage and stem score calculations for WT and *mta* sites, calculated with ViennaRNA using a 150 nt window. Controls were created by taking random windows in TuMV-GFP genome. (E) GC percentage, in a 150 nt window, for each of WT k-mer classes. (F) Percentage of predicted structured sites, calculated with ViennaRNA for each different k-mer class in WT. (G) Density plot showing, in shaded color, the general distribution of m^6^A methylation sites for each k-mer class and each line. In solid the position of all the high confidence predicted secondary structures. The central position of the *m1-m5* sites is indicated by arrows. (H) Graphical representation of the best scoring secondary structures, for each k-mer class and for each line. Figures were generated with RNAfold.

Furthermore, the spatial distribution of these k-mers along the vRNA differed significantly (Fig 5C). We searched for auxiliary upstream or downstream consensus sequences within a 200 nt radius using HOMER and MEME; however, no consistent elements were found across all k-mer categories or within specific groups. We also included host transcripts in this analysis for DRACH/RRACH motifs but failed to identify conserved flanking sequences beyond the central methylation site.

Our previous data demonstrated that a substantial portion of vRNA methylation occurs in the nucleus during early infection stages before equilibration with the cytoplasm. This could result from the nuclear export of methylated vRNA or the activity of a cytoplasmic m^6^A methyltransferase. For this purpose, we investigated FIONA1^93–96^, a predominantly nuclear methyltransferase responsible for U6 snRNA methylation, but found no FIONA1 consensus occurrences within a 10 nt window of our sites (Fig S4C).

We then investigated the role of RNA secondary structure. Initial analysis of a 150 nt window around mapped sites showed that the average GC content in WT was similar to randomized controls, while *mta* showed a slight, non-significant increase (Fig 5D). Using ViennaRNA to calculate a stem score based on Minimum Free Energy (MFE)^97,98^, we found no significant differences between WT, *mta*, and controls (Fig 5D). However, narrowing the analysis to individual k-mer classes in WT revealed that while “GGAU” sites exhibited increased GC content, “Other” and “DRACH” types did not (Fig 5E). Interestingly, structural predictions indicated that many “Other” and “DRACH” k-mers reside in open loop structures, whereas most “GGAU” k-mers are located within hairpins or secondary structures (Fig 5F).

To refine these observations, we analyzed smaller 51 nt windows. Mapping the most highly structured k-mers against the general modification density in the TuMV-GFP genome revealed that “DRACH” and “GGAU” k-mers coincide with the highest density peaks in WT (Fig 5G). Crucially, “Other” k-mers appeared to form a “consortium” surrounding these structured “DRACH” and “GGAU” sites. In *mta*, this organized distribution was lost, being more random and concentrated toward the 3′UTR (Fig 5G).

Structural modelling of these hairpins using RNAfold^98,99^ revealed that while “GGAU” k-mers scored similarly in both lines, “DRACH” and “Other” sites differed. WT “DRACH” consensuses tended to be located in open loops within the stem, whereas those in *mta* were predominantly in smaller loops, similar to “GGAU” sites. “Other” consensuses were primarily located in structured areas of the stem, potentially affecting their biochemical accessibility (Fig 5H).

Finally, we selected five strategic sites (*m1–m5*) characterized by high ONT probability scores and favourable loop structures to test our assumptions (Fig 5G). These were cloned into the *pUBN-GFP* (*pUBN*) expression vector in two window sizes: 51 nt (small, *-s*) and 105 nt (big, *-b*), maintaining the viral reading frame (Fig 6A). Following transient expression in *Nicotiana* leaves, we normalized Western blot fold-changes (FC) by transgene expression levels, consistent with our findings that methylation does not alter transcript stability (Fig 6B-E). While all constructs performed better than the WT, normalization revealed that sites *m4* and *m5* were the most effective (Fig 6F). Notably, expanding the window size generally decreased the normalized FC, except in *m3*, where the larger window included an additional WT-mapped “DRACH” consensus. Similarly, when *m1* and *m2* were fused due to their proximity, the loss of the “DRACH” consensus in *m2* reduced the normalized FC to levels similar to *m1*, whereas *m5*, centered on a stable “DRACH” site, maintained its high normalized FC.

**Figure 6:**
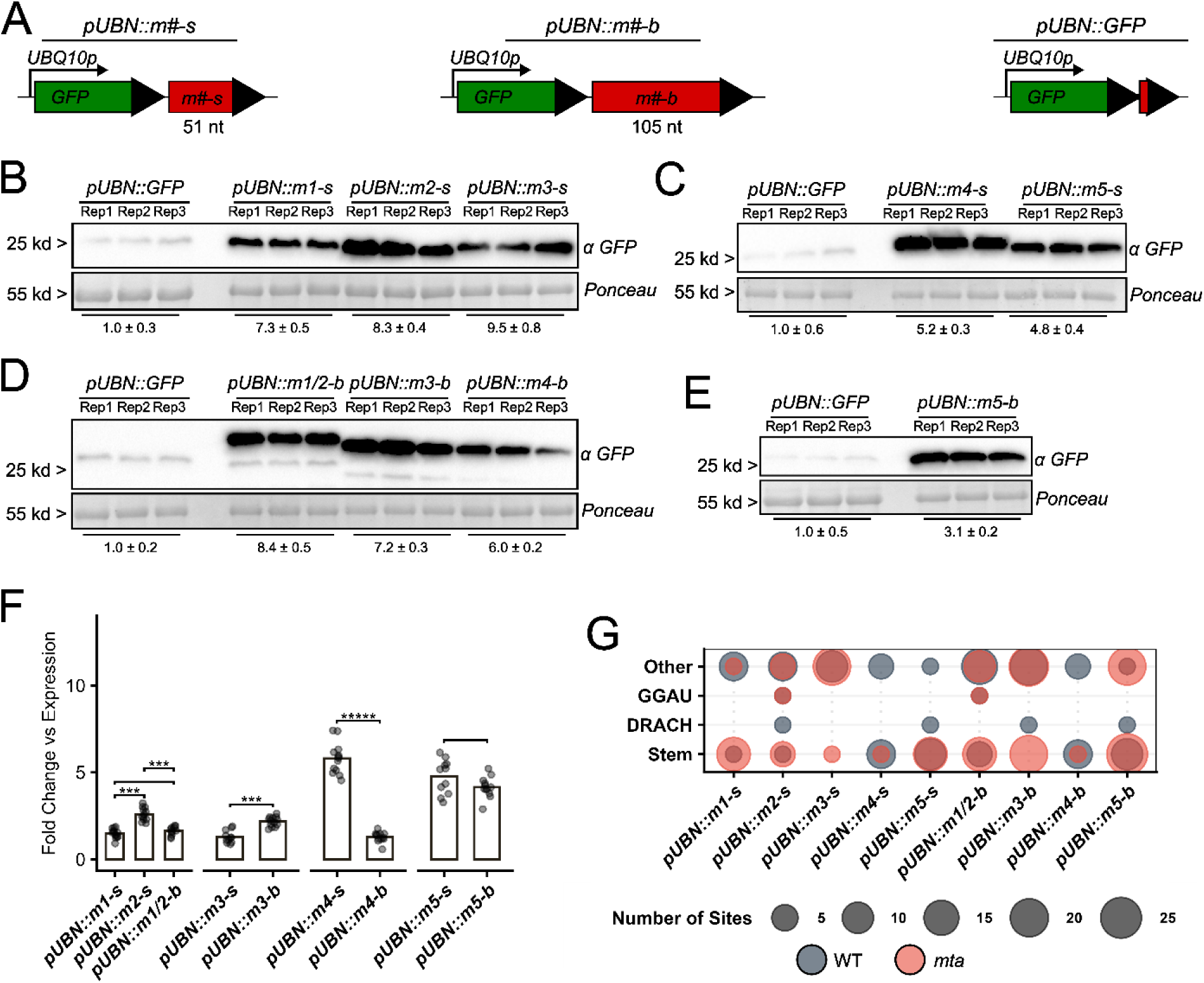
DRACH consensus and the presence of secondary structures favors transgene protein synthesis. (A) Schematic representation of the expression system used in the study. *pUBN::GFP* was obtained by inserting a short oligo containing a stop codon in the cloning site. (B) Western blot analysis of *pUBN::m1-s* to *pUBN::m3-s* transfected leaves 2 days post agro-infiltration. (C) Western blot analysis of *pUBN::m4-s* and *pUBN::m5-s* transfected leaves 2 days post agro-infiltration. (D) Western blot analysis of *pUBN::m1/2-b* to *pUBN::m4-b* transfected leaves 2 days post agro-infiltration. (E) Western blot analysis of *pUBN::m5-b* transfected leaves 2 days post agro-infiltration. We used α-GFP antibodies to identify transgenes. Band intensities were quantified relative to the most intense WT band, with values representing fold changes. Each band corresponds to a biological replicate, with each replicate consisting of four plants, 2 leaves were infiltrated in each plant. (F) RT-qPCR normalization of the protein Fold change, of each expression vector. Data was further normalized by GFP normalized FC. Statistical comparisons between specified groups were performed using two tailed Student’s t-tests. Significance levels are indicated by asterisks (“*”), where the number of asterisks corresponds to -log_10_(p-value), with a minimum threshold set at p-value < 0.05. Non-significant comparisons are not labeled. (G) Map representing the different features of each expression vector, in the stem line the dot represents the number of modified bases hits in a stem predicted area.

These data suggest that TuMV protein expression is not governed by isolated methylation events but by a complex interplay between canonical DRACH sites and surrounding non-canonical clusters, whose functionality is strictly dependent on their secondary structure context.

### Spatially coordinated m^5^C and m^6^A modifications form highly structured epitranscriptomic clusters

While our data suggest a direct relationship between DRACH and “Other” consensus motifs in WT, the previous failure to identify DRACH motifs using HOMER and MEME indicated that a critical detail regarding the formation of these methylation clusters was missing. We suspected that RNA modifications beyond m^6^A may play a role and therefore investigated the contribution of another key modification, 5-methylcytosine (m^5^C). For this, we investigated the m^5^C modification landscape of TuMV-GFP vRNAs using our ONT dataset. Following normalization for read depth and biological replicates, only sites in the upper median were retained for further analysis.

As expected, no global consensus for m^5^C methylation was identified. Sequence-based hierarchical clustering revealed that nearly all m^5^C sites in both WT and *mta* grouped into Cluster 1, which lacks a clear motif. Notably, *mta* mutants possessed only 17% of the m^5^C sites found in the WT. Although m^5^C methylations are primarily catalyzed by the NSUN protein family and do not fundamentally depend on the m^6^A writer complex, the m^5^C distribution in WT strictly mirrored that of m^6^A. This spatial coordination was significantly less prominent in the *mta* mutant. Furthermore, WT m^5^C Cluster 2 specifically resembled a canonical DRACH motif, suggesting a potential overlap in targeting.

We then mapped the distance between each m^6^A k-mer class and m^5^C sites. In WT, the majority of m^5^C methylations were located within 10 nt of an m^6^A modification, a spatial proximity that was lost in *mta*. A significant number of m^5^C sites were specifically clustered near DRACH m^6^A consensuses. Correlation tests between m^6^A and m^5^C residues yielded an R-score of 0.113 for WT compared to -0.258 for *mta*, indicating a higher degree of modification correlation in the WT background.

To refine these findings, we narrowed the correlation window to 20 nt and analyzed the association between m^5^C clusters and m^6^A k-mer classes. While most m^5^C sites associated with the “Other” m^6^A k-mer class in both lines, we observed a substantial enrichment of Cluster 2 m^5^C modifications within DRACH m^6^A consensuses in WT, an effect that was averaged out in *mta*. Remarkably, 92.3% of DRACH sites in WT are located in close proximity to m^5^C modifications. Most significantly, every DRACH consensus located within a secondary structure was found to be adjacent to an m^5^C site. These results indicate that m^6^A and m^5^C modifications are not independent events but rather form highly coordinated epitranscriptomic clusters, where m^5^C potentially acts as a structural stabilizer or co-factor for m^6^A deposition around canonical DRACH motifs.

## Discussion and Conclusion

RNA modifications, particularly N^6^-methyladenosine (m^6^A), play a fundamental role in regulating RNA metabolism across eukaryotic systems. In this study, we investigated the interaction between Turnip Mosaic Virus (TuMV), a widespread +ssRNA potyvirus, and the host methylation machinery in *Arabidopsis*. Our results demonstrate that m^6^A and m^5^C methylation significantly influence TuMV polyprotein synthesis and virulence. Most notably, we identified a novel landscape of non-canonical m^6^A methylation that plays a critical role in regulating viral metabolism.

While many m^6^A-virus interactions negatively impact viral fitness, m^6^A has been shown to enhance replication in some instances, such as Wheat Yellow Mosaic Virus (WYMV)^67,100^. Our findings reveal that the host m^6^A writer complex is essential for supporting TuMV propagation. Using Arabidopsis *mta* mutants, we demonstrated that TuMV infection is significantly impaired in the absence of a functional MTA methyltransferase (Fig 1C, 1D, 1F). In *mta* plants, viral protein expression declined post-infection and failed to recover, indicating that TuMV requires the host methylation machinery to establish a successful infection cycle (Fig 1A, 1D, 1F). Notably, the loss of MTA did not prevent initial infection but severely restricted replication, resulting in viral resilience rather than complete resistance (Fig 2B, 2D–2H).

*Sha et al.*^68^ reported contrasting results in *B. juncea*, where TuMV was negatively regulated by m^6^A. However, the field of plant epitranscriptomics is relatively new, and host-virus interactions are notoriously specific. By expanding our study beyond the writer complex to include reader and erasers, we provided a more comprehensive spectrum of this interaction. While the writer complex is primarily associated with m^6^A, ALKBH*-like* demethylases can target both m^6^A and m^5^C, suggesting that these pathways may be more intertwined than previously thought^33^ (Fig 2D-I). Our *ect* mutant data further clarifies that viral protein expression is heavily regulated by reader proteins during early infection stages, affecting both protein levels and vRNA titers (Fig S2B-S2G). Interestingly, the dependency on host ECTs does not appear to confer host resilience in the long term, as the viral infection was not suppressed in any of the examined *ect* mutants. *Martinez-Perez et al.*^50^ demonstrated that the introduction of mutated *ECT* genes was sufficient to restore AMV virulence in an *alkbh9b* background. Although our TuMV model behaves differently from AMV, we observed similar effects within two distinct ECT groups. Specifically, the *ect2/3/4* and *ect4* mutant showed increased viral protein accumulation compared to the WT, indicating a potential antagonistic role in regulating TuMV protein synthesis (Fig S2B-S2F). Conversely, we observed the opposite scenario with *ect5*, *ect7*, and *ect9* (Fig S2B-S2E, S2G). While we primarily focused on *ect4* and *ect9*, exploring higher-order *ect* mutant lines will be necessary to further clarify these roles. These findings highlight a critical aspect of virus-epitranscriptome interactions, challenging the notion of an absolute beneficial or antagonistic role of host modifications on viral fitness. Our analysis shows that high levels of vRNA methylation occur in the nucleus, where the writer complex is localized (Fig 4A-C). *Feng et al.* demonstrated that the CMV2b protein promotes the relocalization of MTA into nuclear condensates, inhibiting its functions^101^. While some viruses who benefit from host methylation relocate the writer complex to the cytoplasm^67^, we demonstrated that TuMV-GFP vRNA is present in the host nucleus and is likely targeted by the writer complex there. Although we detected traces of viral protein in the nucleus, proteins such as NIb and NIa-pro, which are within the observed molecular weight, are known to localize to the nucleus during potyviral infections^102–105^. Our data demonstrates that the viral RNA itself is also present in the nuclear fraction, suggesting that these proteins may be co-localized with vRNA to facilitate nuclear-specific processes such as m^6^A and m^5^C modification (Fig 4D).Potyviral CP, NIb, and NIa proteins have been previously documented to play active roles in either favoring or inhibiting viral methylation^44,68,68,106,107^. This is achieved through the recruitment of host demethylases or by interacting with the writer complex and accessory factors to suppress vRNA methylation^44,68,106,107^. Conversely, some viruses utilize viral proteins to recruit methyltransferases, thereby promoting vRNA methylation to enhance their fitness^67,100^. For instance, *Zhang et al.* demonstrated that a single polymorphism in the *TaMTB* gene is sufficient to disrupt the interaction between the WYMV NIb protein and the host MTB subunit^67^. This underscores the constant evolutionary pressure on both viruses and their hosts to develop mechanisms for evading defenses or hijacking host metabolism. Consequently, even closely related viruses within the same family, sharing similar genomic architectures, can exhibit markedly different strategies to deal with the host epitranscriptome. To our knowledge, little to no explanation has been provided to justify these divergent behaviours. Additionally, traditional m^6^A immunoprecipitation-based techniques (MeRIP-seq) are prone to including false positives^108^. To overcome these limitations, we combined our MeRIP data with Oxford Nanopore Technologies (ONT) direct RNA sequencing, which allowed us to obtain an in-depth, comprehensive understanding of the viral epitranscriptomic status.

Our MeRIP-sequencing profile revealed three primary peaks localized near the 3′UTR of the virus (Fig 3A). These peaks increased in intensity toward the 3′UTR, with the final peak exhibiting the strongest signal. The status of this terminal peak as a major methylation target was further confirmed through the analysis of vRNA isolated from *MTA-YFP* plants, which function as a mild overexpressor (Fig 3A). Interestingly, while our MeRIP-sequencing profile shares some similarities with the findings of *Sha et al.*^68^, it displays a distinct architecture. Specifically, we observed the highest intensity in the final peak, whereas their strongest signal was localized near our first and second peaks (Fig 3A). The application of ONT direct RNA sequencing further clarified these observations by revealing the exact coordinate of each modified base at single-nucleotide resolution (Fig 3B).

However, the most intriguing discovery did not come from TuMV-GFP vRNA itself but rather from *Arabidopsis* transcripts. Interestingly, while *mta* mutants maintain a methylation pattern similar to WT for host mRNAs (Fig 3E), the viral methylation pattern is severely compromised; specifically, the canonical DRACH motifs are reduced and misplaced (Fig 3B, 3C). Conversely, “GGAU” consensuses are enhanced in *mta*, suggesting the activity of a secondary methyltransferase that may compensate for *MTA* or act independently based on RNA secondary structure (Fig 3C).

Previous investigations into virus-epitranscriptome interactions have focused primarily on the presence or absence of methylation on viral transcripts and how these modifications either favor or inhibit infection. We have expanded on this by identifying and classifying distinct methylation classes (Fig 3D). We propose that the “Other” non-canonical consensuses, which we identified exclusively in viral transcripts, serve as a specialized regulatory mechanism. Notably, these modifications not only yielded the highest ONT scores but also closely mirrored our MeRIP-sequencing profile, providing robust cross-platform validation for their existence and distribution. Our stability assays indicate that these modifications primarily influence translation efficiency rather than RNA degradation (Fig 4E,4F). We also observed that these “Other” non-canonical methylations cluster around structured sites containing DRACH and GGAU consensuses in the WT, whereas they cluster around GGAU consensuses only in *mta,* albeit structured DRACH consensuses being present (Fig 3G). Secondary structures have been reported in several viral genomes and are essential for various aspects of viral metabolism, including replication and translation initiation^109–111^. Conversely, such highly structured regions can serve as hallmarks for viral recognition by the host immune system, potentially acting as a liability for the virus^112–114^.

*Arribas-Hernández et al.*^91^ previously reported that in Arabidopsis ECT2 prefers to bind to DRACH sites flanked by U-rich motifs. By analogy, one might expect other ECT proteins to exhibit similar binding preferences; however, we could not identify conserved motifs around any of the k-mer classes identified in the viral genome. Furthermore, the “Other” methylation class lacked a conserved consensus even after hierarchical clustering. Conversely, our data indicates that ECT4 and ECT9 have opposing roles in regulating viral protein expression (Fig S2B-S2G). ECT4 has been previously functionally correlated with ECT2 and ECT3^75–77^, and our results support this relationship, as *ect4* and *ect234* mutants exhibited the most dramatic fold change increase in viral proteins compared to WT(Fig S2B-S2F). However, the lack of a conserved sequence preference for ECT4 or ECT9 suggests that their recruitment to viral transcripts may be governed by factors other than primary sequence, such as RNA secondary structure or nearby modifications^75,115^. Beyond acting as targets for YTH domain-containing proteins, m^6^A modifications possess the ability to stabilize single-stranded RNA regions, thereby altering the overall RNA secondary structure^116–118^. Interestingly, most methyltransferases exhibit a preference for unstructured motifs, as the majority of previously mapped m^6^A sites occur in such regions^116,119,120^. This observation holds true for our dataset: with the exception of GGAU motifs, both DRACH and “Other” consensuses are primarily located in unstructured regions, specifically, regions with high Minimum Free Energy (MFE) (Fig 5E). Notably, structured methylated regions, which account for approximately 32% of the mapped methylations (39% in WT and 26% in *mta*), tend to cluster together in WT, whereas this clustering is abolished in the *mta* mutant (Fig 5F, 5G).

Synthesizing these observations, we found that despite lacking distal consensus sequences, the TuMV methylation pattern is spatially distributed around structured DRACH and GGAU motifs. In WT plants, this pattern extends to neighbouring structured “Other” sites, forming distinct “methylation islands.” We propose a seeding model wherein an MTA-containing writer complex, likely in association with a secondary methyltransferase governed by structural rather than sequence-based preferences, recognizes and deposits strategic m^6^A marks to establish a functional epitranscriptomic consortium. In *mta* the lack of a functional writer complex makes this secondary methyltransferase becomes the primary enzyme, highlighting the higher concentrations of GGAU sites (Fig 3C).

Furthermore, our transgene expression experiments confirmed that high protein expression requires the combination of a canonical DRACH consensus within or near a structured RNA region (Fig 6F, 6G). This is clearly illustrated by the *m1* and *m2* expression constructs, where the normalized protein levels were reduced upon the loss of the DRACH consensus (Fig 6B, 6D, 6F). One outlier was the *m4* site, which exhibited high normalized FC despite the absence of DRACH or GGAU motifs. We hypothesize that a DRACH-like consensus may have been inadvertently created during the cloning process; indeed, upon extending the cloned insert, the normalized FC dropped. In contrast, the *m5* construct (centered on a DRACH site) remained stable, and the *m3* construct showed increased normalized FC upon the addition of a DRACH motif (Fig 6B-6F).

While this model explains the reduced protein levels observed in writer mutants, and the eventual viral demise, the “Other” m^6^A spectrum remains a point of interest, as it extends far beyond these structured methylation islands in both *mta* and WT, lacking identifiable structural or sequence features beyond the methylated adenosine itself (Fig 3B-3D, 5G). To explain the broader m^6^A spectrum, we investigated the m^5^C landscape. In plants, m^5^C is primarily catalyzed by NSUN*-like* methyltransferases, which lack a clear consensus motif but typically exhibit a preference for GC-rich regions^121,122^. Interestingly, the m^5^C sites in our dataset did not show significant GC enrichment compared to the control (Fig S4D). However, we observed that while underrepresented, m^5^C Cluster 2 contained a DRACH consensus in the WT (Fig 7A–7C). Additionally, the majority of m^5^C modifications in the WT were located within 20 nucleotides of an m^6^A site, a spatial coordination that was abolished in the *mta* mutant (Fig 7D, 7E). Notably, while DRACH motifs account for only 5% of total m^6^A sites, they represent 33.3% of the m^6^A modifications surrounding m^5^C sites in Cluster 2. The high frequency of DRACH motifs in the immediate vicinity of m^5^C sites suggests that this spatial arrangement is non-stochastic.

**Figure 7:**
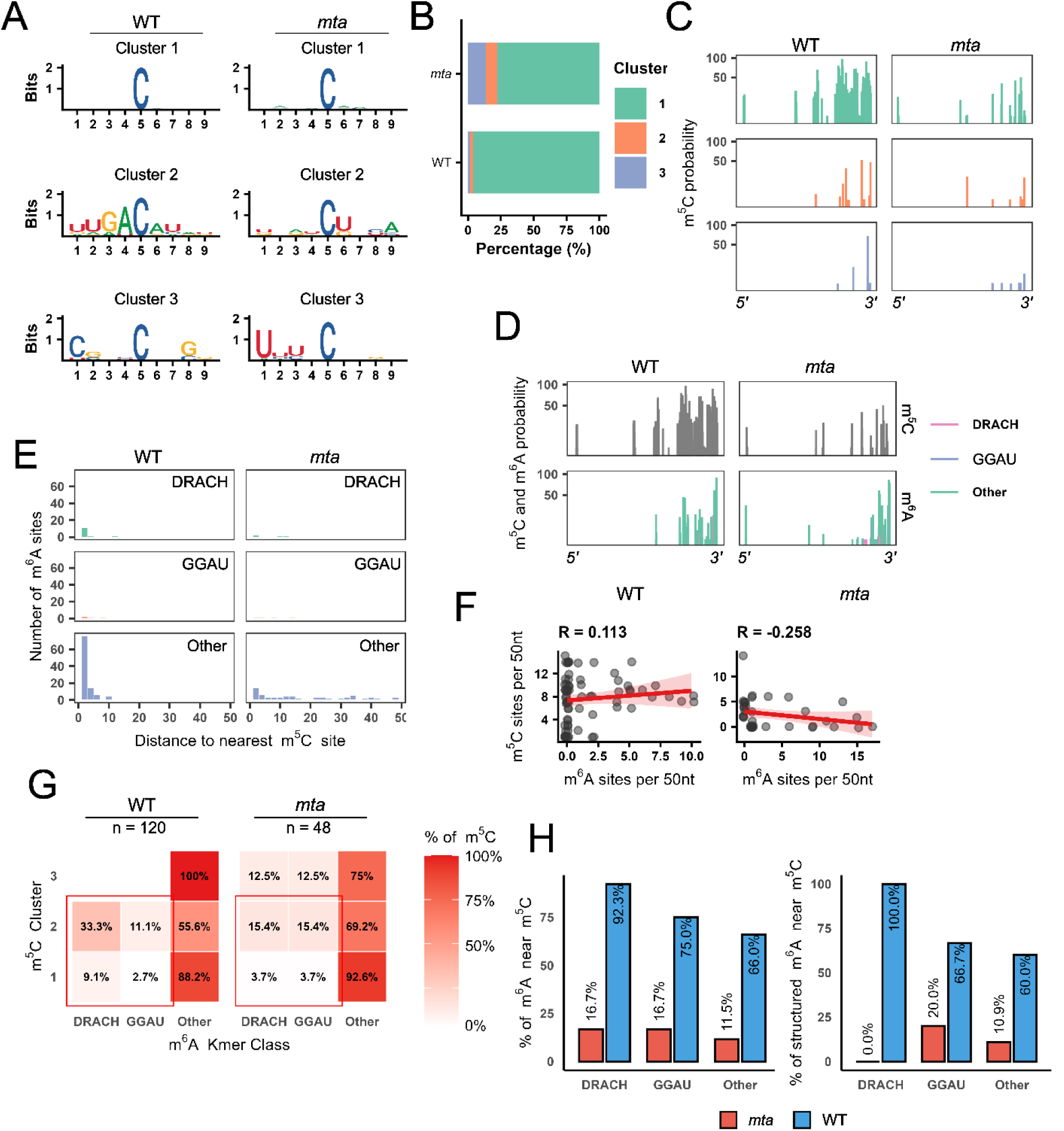
m^5^C modifications cluster around DRACH modifications in structured areas of the viral genome. (A) Nucleotide logos representing the three consensus clusters identified in the ONT analysis. Each logo was calculated independently for each plant plant line. (B) Bar plot showing the different percentages of m^5^C clusters in each plant line. (C) Spatial distribution of m^5^C clusters in each plant line. (D) Comparison of the spatial distribution of m^5^C and m6A in each line. (E) Bar plot showing the average proximity of m5C modifications to each m^6^A consensus. (F) Correlation plot between m^5^C and m^6^A sites within a 50 nt window (G) Heatmap showing the percentage of modifications from each m^5^C cluster located within 20 nt of an m^6^A consensus for each line. Relative percentages were calculated by independent normalization for each plant line. (H) Bar plot showing the percentages of each m6A consensus that are within 20 nt from a m5C modification (left) and m6A consensus located in structured ares that are within 20 nt from a m5C modification (right). Relative percentages were calculated by independent normalization for each plant line.

**Figure 8:**
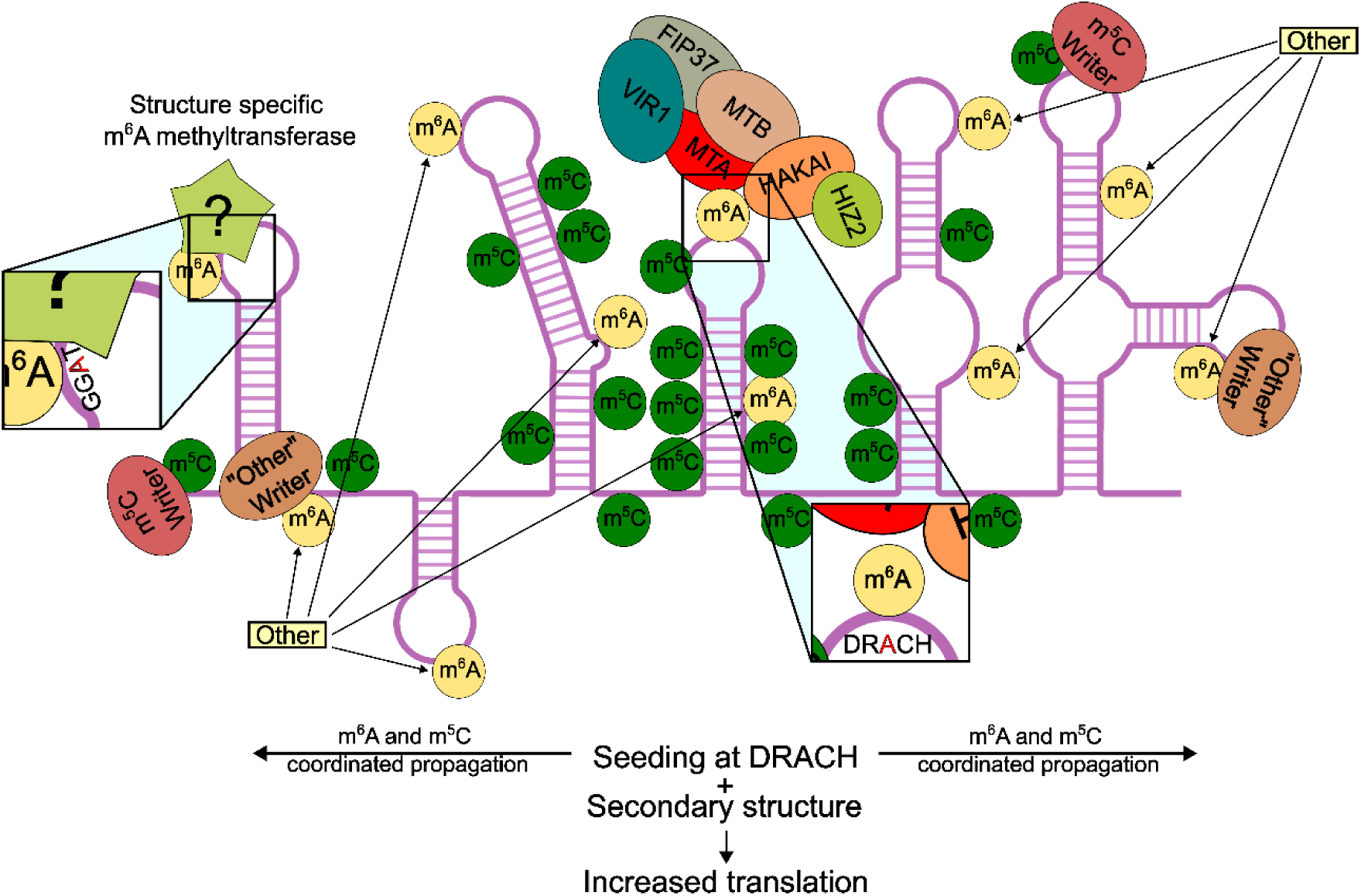
Proposed model of m^6^A and m^5^C-mediated regulation of TuMV infection. This schematic model illustrates the fate of TuMV-GFP infection based on host methylation status. The initial seeding necessitates a fully functioning writer complex that then generates a cascade of events that ends in methylation islands which have a major impact in regulating vRNA protein synthesis.

Although we currently lack experimental validation using NSUN mutants (such as *trm4b*), a functional link between m^5^C and m^6^A has recently been established in animal models. For instance, *Orji et al.* demonstrated the co-localization of m^6^A-methylated RNAs and m^5^C reader proteins *in vitro*, while *Li et al.* reported a mutualistic co-regulatory effect of m^6^A and m^5^C on *p21* transcripts^123,124^. Furthermore, m^5^C has emerged as a significant player in viral infections across both animal and plant systems, mirroring the importance of m^6^A^24,55,56^.

Finally, we observed a striking spatial correlation between m^6^A and m^5^C (Fig 7D–F). In WT plants, 92.3% of DRACH sites are located in close proximity to m^5^C modifications, specifically within secondary structures (100%), a coordination that is lost in the *mta* mutant. This suggests that m^6^A and m^5^C do not function in isolation but work in tandem to establish a functional “epitranscriptomic landscape” that optimizes viral fitness (Fig 7G,7H).

## Conclusion

In conclusion, this study redefines our understanding of the potyvirus life cycle by uncovering a critical nuclear phase during which TuMV hijacks the host’s m^6^A and m^5^C methylation pathways. We propose a “seeding” model of viral methylation, wherein the canonical writer complex deposits m^6^A at primary DRACH motifs to serve as anchor points. This initial event likely triggers a cascade of secondary m^6^A and m^5^C modifications, guided more by RNA secondary structure than primary sequence, to form a coordinated epitranscriptomic consortium.

In the absence of this nuclear priming, the virus exhibits an aberrant methylation landscape that fails to support efficient polyprotein synthesis, leading to a loss of viral fitness. While it remains to be determined whether these methylation patterns primarily influence host DICER-mediated defense pathways by altering RNA topology or function primarily to recruit specific ECT readers, our data strongly support the latter. Furthermore, whether this strategy enables the virus to selectively recruit beneficial readers or serves to “saturate” the vRNA to evade host immune recognition remains an open question. Nevertheless, these findings highlight the profound complexity of plant-virus interactions and identify the host’s methylation apparatus as a high-potential target for engineering broad-spectrum viral resilience in crops.

### Limitations of the study

While our findings strongly support the critical dependence of TuMV on the host methyltransferase complex, the precise roles of specific ECT proteins in coordinating viral replication and translation remain to be fully elucidated. Investigating higher-order mutant lines combining various writer and reader deficiencies will be essential to clarify these functional nuances. Future research should aim to dissect the molecular mechanisms by which m^6^A modifications influence the TuMV life cycle, specifically determining whether distinct ECT proteins function as viral repressors or enhancers in a context-dependent manner. Furthermore, studies utilizing *trm4b*, *nop2*, and *aly*-family mutants, either alone or in combination with m^6^A mutants, will be necessary to strengthen the functional correlation between m^6^A and m^5^C modifications. This study also does not explore whether TuMV actively modulates host methylation dynamics or how the viral epitranscriptome influences sequential infections or viral synergy. Notably, as *Martinez-Perez et al.* demonstrated, viral interactions with the host methylome can vary significantly even among members of the same viral family^81,81,125–128^. While our study focused primarily on TuMV, challenging *Arabidopsis* mutants with other potyviruses or unrelated viral families could provide a broader understanding of the conserved versus divergent mechanisms governing host epitranscriptome-virus interactions.

## Supporting information

The figure description is already present in the PDF

## Acknowledgements

This work was supported by KAUST funding (BAS/1062-01-01) awarded to HH. The authors extend their gratitude to all lab members for their support. We would like to thank Professor Magdy M. Mahfouz and Dr. Monika Chodasiewicz for generously providing the *p35S::TuMV-GFP* vector and *ect* mutant seeds, respectively. Additionally, we thank Mirko Celii for his assistance with R coding and Anamica Rawaat for helping with confocal microscopy.

## Author Contributions

Conceptualization: A.S., N.S., and H.H.; Methodology: N.S., A.S., and H.H.; Investigation and Data Curation: N.S.; Formal Analysis and Data Processing: K.N. and N.S.; LC/MS Analysis: M.T.; Writing-Original Draft Preparation: N.S.; Writing-Review and Editing: H.H., K.N. and A.S.; Supervision: H.H. and A.S.; Funding Acquisition: H.H. (KAUST BAS/1062-01-01).

## Supplementary data

Supplementary data can be found at the in the Supplementary data pdf.

## Data availability

All sequencing data generated in this study, including RNA-seq, MeRIP-seq, and m6A Oxford Nanopore Technologies (ONT) raw data, have been deposited in a public repository under accession number SUB16132572. Additional data supporting the findings of this study are available from the corresponding author upon reasonable request.

