## Supplementary material for "Turnip mosaic virus co-opts host RNA methylation machinery to orchestrate plant infection": The figure description is already present in the PDF

### Supplementary Figures

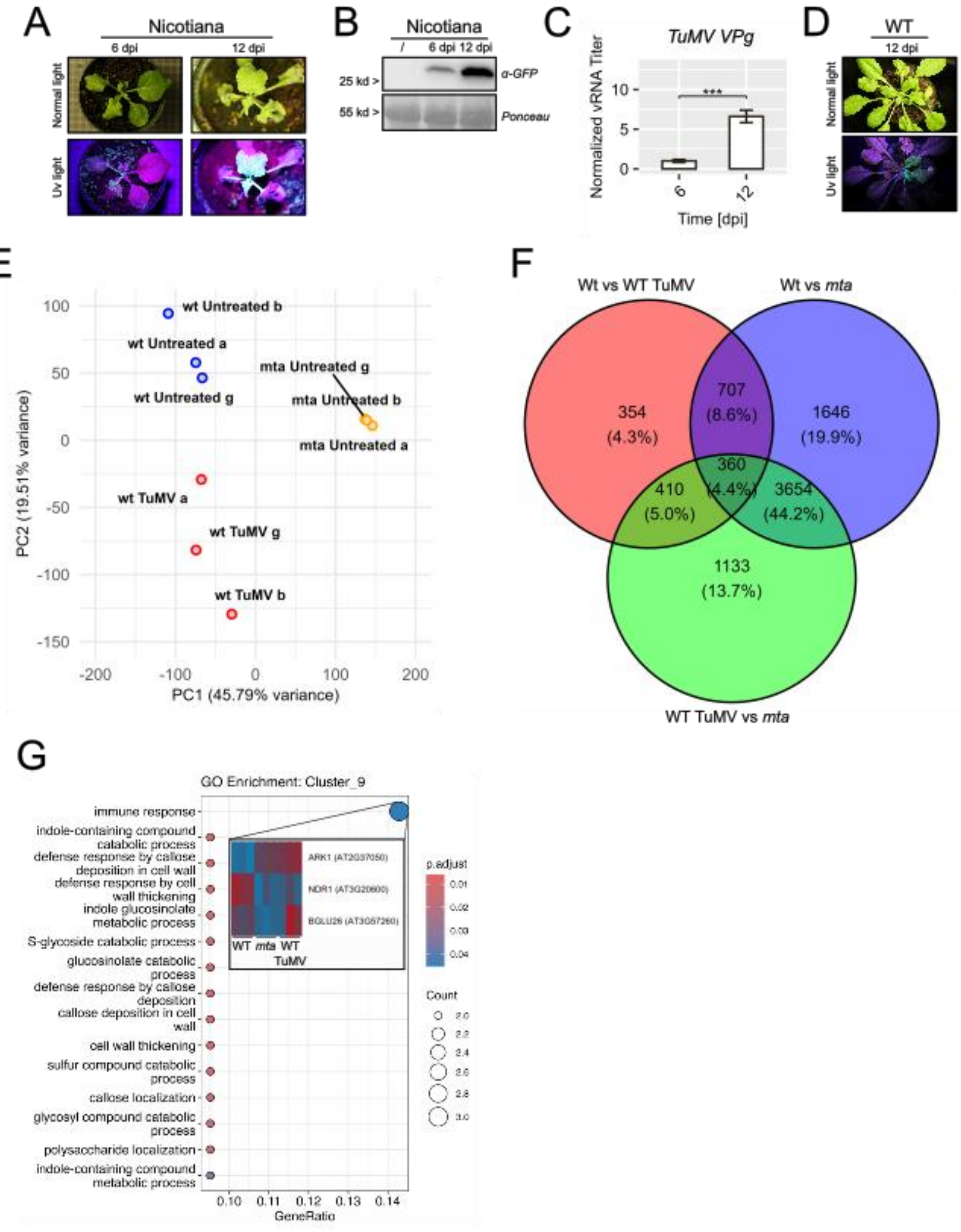


**Figure S1: TuMV-GFP initial testing and transcriptome analysis**

(A) Panel showing Nicotiana, infected with TuMV-GFP at different time points.
(B) Western blot analysis of TuMV-GFP-infected samples using α-GFP antibodies at different time points. Band intensities were quantified relative to the most intense WT band, with values representing fold changes. Each band corresponds to a biological replicate, with each replicate consisting of four plants.
(C) RT-qPCR quantification of the viral titer, using *TuMV VPg* as target, at different time points. Each value represents the mean of six biological replicates, with three technical replicates per biological sample. Error bars represent the variance of each group. Statistical comparisons between specified groups were performed using t-tests. Significance levels are indicated by asterisks ("*"), where the number of asterisks corresponds to $-\log_{10}$(p-value), with a minimum threshold set at p-value < 0.05. Non-significant comparisons are not labeled.

(D) Panel showing TuMV-GFP infected WT plants at 12 dpi.

(E) Principal Component Analysis (PCA) of the transcriptome sequencing data, highlighting significant differences between *mta,* WT untreated and TuMV-GFP-infected samples. PCA was performed to assess overall transcriptomic variation between conditions.

(F) Venn diagram showing the differential expressed genes in *mta,* WT untreated and TuMV-GFP-infected samples.

(G) Gene ontology and heatmap analysis of cluster 9, showing which differentially expressed genes are in the immune response category.

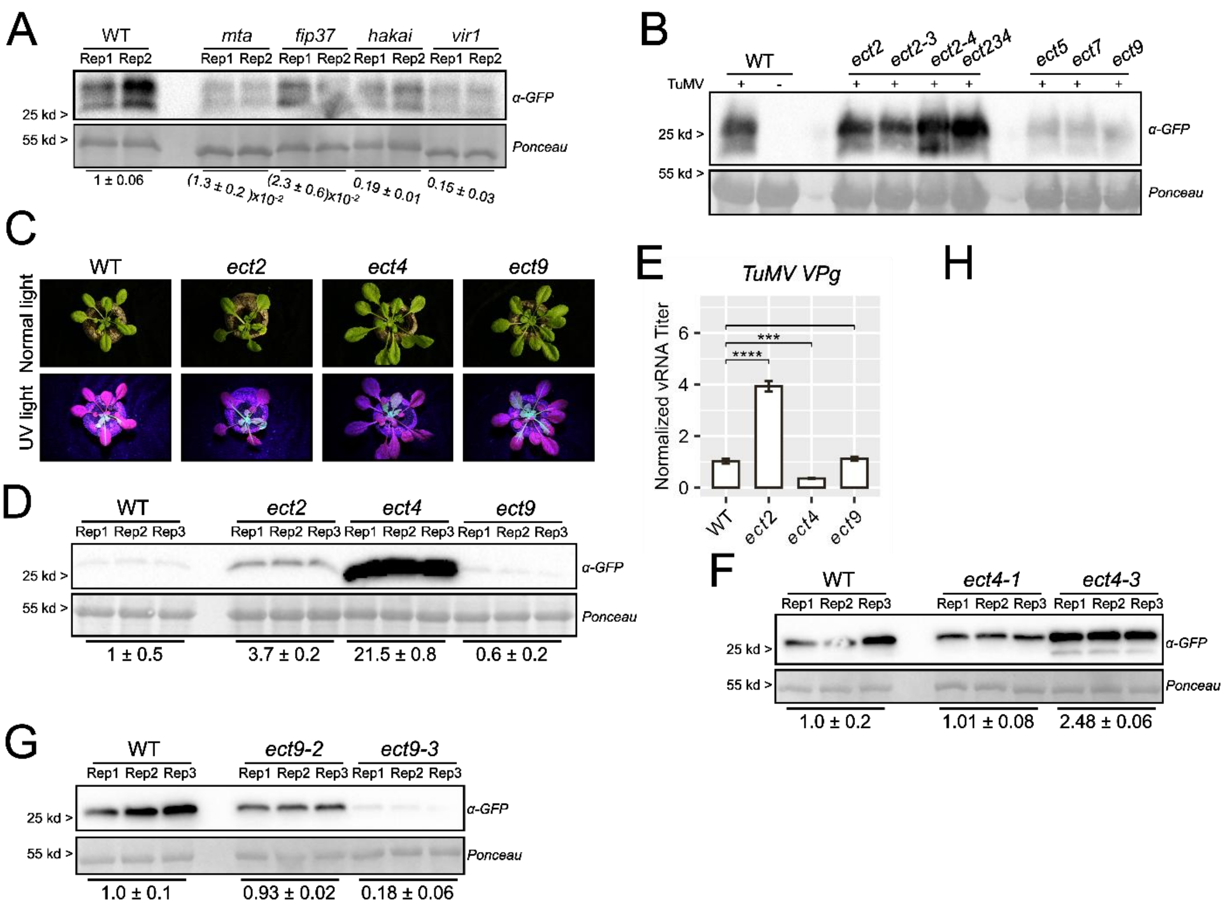


**Figure S2: ECTs impact on TuMV-GFP and RNA methylation and structural data**

(A) Western blot analysis of TuMV-GFP-infected WT and writer mutants using α-GFP antibodies at 1 dpi. Band intensities were quantified relative to the most intense WT band, with values representing fold changes. Each band corresponds to a biological replicate, with each replicate consisting of four plants.

(B) Western blot analysis of TuMV-GFP-infected WT and *ect* mutants using α-GFP antibodies at 2 dpi. Band intensities were quantified relative to the most intense WT band, with values representing fold changes. Each band corresponds to a biological replicate, with each replicate consisting of four plants.

(C) Panel showing TuMV-GFP infected WT and *ect* mutants at 12 dpi.

(D) Western blot analysis of TuMV-GFP-infected WT and *ect* mutants using α-GFP antibodies at 3 dpi. Band intensities were quantified relative to the most intense WT band, with values representing fold changes. Each band corresponds to a biological replicate, with each replicate consisting of four plants.

(E) RT-qPCR quantification of the viral titer, using *TuMV VPg* as target, at 3dpi. Each value represents the mean of six biological replicates, with three technical replicates per biological sample. Error bars represent the variance of each group. Statistical comparisons between specified groups were performed using t-tests. Significance levels are indicated by asterisks ("*"), where the number of asterisks corresponds to $-\log_{10}$(p-value), with a minimum threshold set at p-value < 0.05. Non-significant comparisons are not labeled.

(F) Western blot analysis of TuMV-GFP-infected WT and *ect4* mutant's alleles using α-GFP antibodies at 3 dpi. Band intensities were quantified relative to the most intense WT band, with values representing fold changes. Each band corresponds to a biological replicate, with each replicate consisting of four plants.

(G) Western blot analysis of TuMV-GFP-infected WT and *ect9* mutant's alleles using α-GFP antibodies at 3 dpi. Band intensities were quantified relative to the most intense WT band, with values representing fold changes. Each band corresponds to a biological replicate, with each replicate consisting of four plants.

* - possible false positive

| Rank | Motif | P-value | log P-pvalue | % of Targets | % of Background | STD(Bg STD) |
|---|---|---|---|---|---|---|
| 1 * | 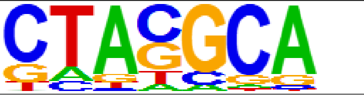  | 1e-3 | -8.870e+00 | 34.09% | 12.23% | 10.4bp (14.3bp) |
| 2 * | 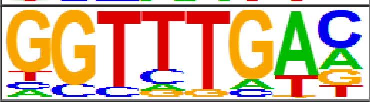  | 1e-3 | -8.080e+00 | 25.00% | 7.49% | 14.4bp (13.3bp) |
| 3 * | 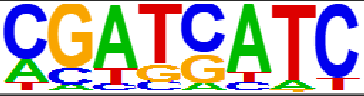  | 1e-3 | -7.686e+00 | 36.36% | 15.18% | 8.3bp (15.0bp) |
| 4 * | 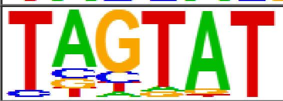  | 1e-3 | -6.909e+00 | 31.82% | 13.04% | 14.0bp (14.7bp) |
| 5 * | 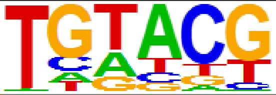  | 1e-2 | -5.336e+00 | 61.36% | 40.85% | 13.3bp (15.9bp) |
| 6 * | 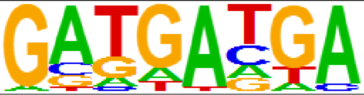  | 1e-2 | -5.325e+00 | 18.18% | 6.15% | 7.9bp (13.7bp) |
| 7 * | 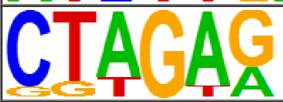  | 1e-2 | -4.772e+00 | 25.00% | 11.30% | 14.2bp (15.6bp) |
| 8 * | 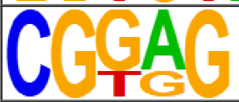  | 1e-1 | -4.523e+00 | 31.82% | 16.85% | 12.7bp (17.7bp) |
| 9 * | 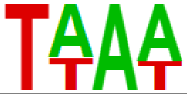  | 1e-1 | -2.626e+00 | 50.00% | 38.12% | 14.8bp (15.5bp) |
| 10 * | 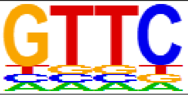  | 1e-1 | -2.502e+00 | 47.73% | 36.42% | 12.8bp (15.8bp) |
| 11 * | 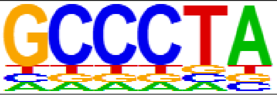  | 1e-1 | -2.438e+00 | 2.27% | 0.22% | 0.0bp (9.7bp) |

**Figure S3: HOMER results on TuMV-GFP MeRIP-sequencing data**

Table summarizing HOMER motif-finding analysis on TuMV-GFP MeRIP-sequencing peaks.

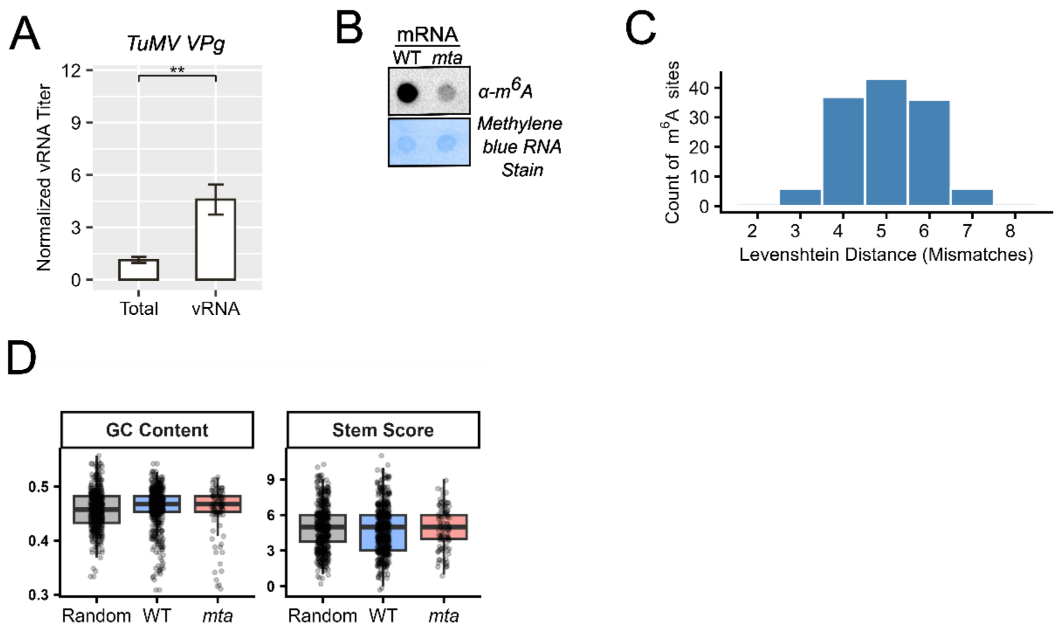


**Figure S4 vRNA purification and other supplementary informations**

(A) RT-qPCR quantification of the viral titer, using *TuMV VPg* as target, comparing total RNA vs viral RNA. Each value represents the mean of six biological replicates, with three technical replicates per biological sample. Error bars represent the variance of each group. Statistical comparisons between specified groups were performed using t-tests. Significance levels are indicated by asterisks ("*"), where the number of asterisks corresponds to $-\log_{10}$(p-value), with a minimum threshold set at $p$-value $< 0.05$. Non-significant comparisons are not labeled.

(B) Dot blot analysis comparing $m^6A$ methylation levels in mRNA isolated from non-infected WT and *mta* plants. Each dot represents equimolar RNA pooled from four independent biological replicates.

(C) Graphical representation of the Levenshtein distance between each mapped $m^6A$ site and the METTL16 consensus motifs

(D) **Average GC content and loop score** of $m^5C$ sites calculated within a 75 nt interval surrounding the mapped methylation sites, comparing WT and *mta* sites. Random 151 nt sequences from the TuMV-GFP genome were utilized as a control.
